# Mechanochemical cues control the coupling of metabolic and migratory patterns in cancer

**DOI:** 10.64898/2026.08.21.746296

**Authors:** Alice Amitrano, Debanik Choudhury, Brent Ifemembi, Alexandros Afthinos, Konstantin Stoletov, Qinling Yuan, Sanjiban Nath, Bishwa Ranjan Si, Bhawana Agarwal, Gianna Graziano, Jack Gao, Anneliese Ceisel, Matthew Hauf, Sean X. Sun, Andrew J. Ewald, Miguel A. Valverde, John D. Lewis, Jeff Mumm, Konstantinos Konstantopoulos

## Abstract

Confined migration is essential for metastasis, yet how cells adapt their migratory and metabolic programs across stiffness-varying microenvironments remains unclear. We uncover a stiffness-dependent mechano-metabolic switch governing migration. In stiff microchannels, cells utilize the osmotic engine model (OEM), relying on NHE1 activity, front-polarization, and glycolysis. In soft microchannels, migration is OEM-independent and requires pyruvate-fueled oxidative phosphorylation (OxPHOS). This OxPHOS-driven motility depends on Arp3, β1-integrin and integrin-linked kinase, which increase membrane tension in confinement that in turn triggers TRPM7-mediated calcium influx and RhoA-/myosin-II contractility. Activating and polarizing NHE1, via overexpression, hypoxia or elevated viscosity, restore OEM- and glycolysis-dependent migration in soft microchannels, bypassing the need for actin polymerization *in vitro* and in chick embryos. Mitochondria addition reinstates Arp3 polarization and enhances migration in NHE1-overexpressing cells, enabling engagement of both mechanisms *in vitro* and in zebrafish. These findings uncover a previously unrecognized mechano-metabolic link, revealing that intracellular rewiring overrides stiffness-dependent metabolic demands.

---

The physical properties of the tumor microenvironment, including confinement^1^ and stiffness^2,3^, critically regulate cancer cell proliferation, migration, and differentiation. During metastasis, cells navigate through narrow three-dimensional (3D) tracks, either pre-existing or generated by extracellular matrix (ECM) remodeling, requiring them to dynamically adapt their migratory strategies^1,4,5^. Advanced *in vitro* platforms that recapitulate the interplay between confinement and stiffness have been essential for uncovering how cells integrate these cues to navigate complex environments^6^.

In stiff confining channels, migration can occur independently of actin, myosin II, or β1-integrin, via the Osmotic Engine Model (OEM), which relies on polarized distribution of ion transporters and ion channels, such as NHE1 and SWELL1, which coordinate cycles of isosmotic swelling and shrinkage at the cell front and rear, respectively^7,8^. Disruption of this polarization pattern impairs migration in stiff microchannels, underscoring the role of ion transport dynamics in confined motility^8^. Despite extensive studies in stiff microchannels, the mechanisms by which cells adapt their migratory strategies to substrates of differing stiffness remain poorly understood.

Another critical yet less explored aspect of this adaptation is cellular metabolism, which enables cancer cells to couple energy production with the demands of migration^5,9,10^. Metabolic reprogramming, characterized by shifts between glycolysis and mitochondrial oxidative phosphorylation (OxPHOS), is a hallmark of cancer progression and plays a key role in supporting metastasis^11,12^. However, findings on the mechanical regulation of cancer cell metabolism and motility across substrates of varying stiffness remain context dependent. Stiff matrices promote glycolysis in hepatocellular carcinoma^13^ and lung cancer cells^14^, while glioblastoma cells retain a glycolytic phenotype in soft, brain-mimetic environments^15^. In contrast, pancreatic ductal adenocarcinoma^16^ and breast cancer cells^17^ can engage mitochondrial metabolism in response to increased matrix stiffness to support invasive behavior. These discrepancies suggest that the cell metabolic adaptation may not be universally dependent on substrate stiffness. Instead, we herein demonstrate that cell intracellular rewiring from actomyosin- to OEM-based motility can override the influence of matrix stiffness on cellular bioenergetic requirements.

## The roles of OEM and the cytoskeleton in cell motility within stiff and soft microchannels

To investigate how substrate stiffness influences confined cancer cell migration, we fabricated microfluidic devices containing an array of parallel, straight microchannels (Width x Height = ∼3 µm × 10 µm) using either polydimethylsiloxane (PDMS; >1,000 kPa)^7,8^ or polyacrylamide gels^3^ of prescribed stiffness (7, 15 or 35 kPa). Given the critical role of the OEM in driving migration under confinement inside stiff microchannels^7,8^, we first examined whether its key regulator, the Na⁺/H⁺ exchanger NHE1, exhibits stiffness-dependent polarization. While NHE1 is preferentially enriched at the leading edge of breast cancer MDA-MB-231 cells inside stiffer microchannels, including PDMS-based and 35 kPa PA gel-based, it fails to polarize in softer 15 kPa microchannels (**Fig. 1a,b**). A similar trend is observed in human osteosarcoma HOS cells (**Extended Data Fig. 1a,b**), as well as triple negative breast cancer patient-derived xenografts (PDX) cells (**Extended Data Fig. 1c,d**), where NHE1 polarized in PDMS-based but not in 15 kPa polyacrylamide-based microchannels. Moreover, NHE1 activity also displays a stiffness-dependent behavior, gradually diminishing as stiffness decreases (**Fig. 1c**). Reduced NHE1 polarization and activity are also observed in cells within even softer (7 kPa) microenvironments (**Extended Data Fig. 1e-g**). In agreement with these findings and prior work^7,8^, NHE1 knockdown^18^ suppresses migration in stiffer (35 and >1,000 kPa) but not softer (15 kPa) microchannels (**Fig. 1d**). In accord with the OEM^7,8^, AQP5 knockdown^18^ reduces confined migration in 1,000 kPa, while migration in 15 kPa microchannels remains unaffected (**Extended Data Fig. 1h**). Consistent with the role of Yes-Associated Protein (YAP) as a stiffness mechanotransducer^19,20^, whose nuclear localization is reduced in soft microenvironments (**Fig. 1e,f**), YAP inhibition (**Extended Data Fig. 1i**) disrupts NHE1 activity and leading-edge polarization in stiff microchannels (**Extended Data Fig. 1j-l**), recapitulating the responses observed in control cells within soft microchannels. These findings support a model in which YAP-dependent NHE1 polarization and activity are critical to migration in stiff microchannels but become dispensable in soft ones (**Fig. 1a-c, g**).

**Figure 1.**
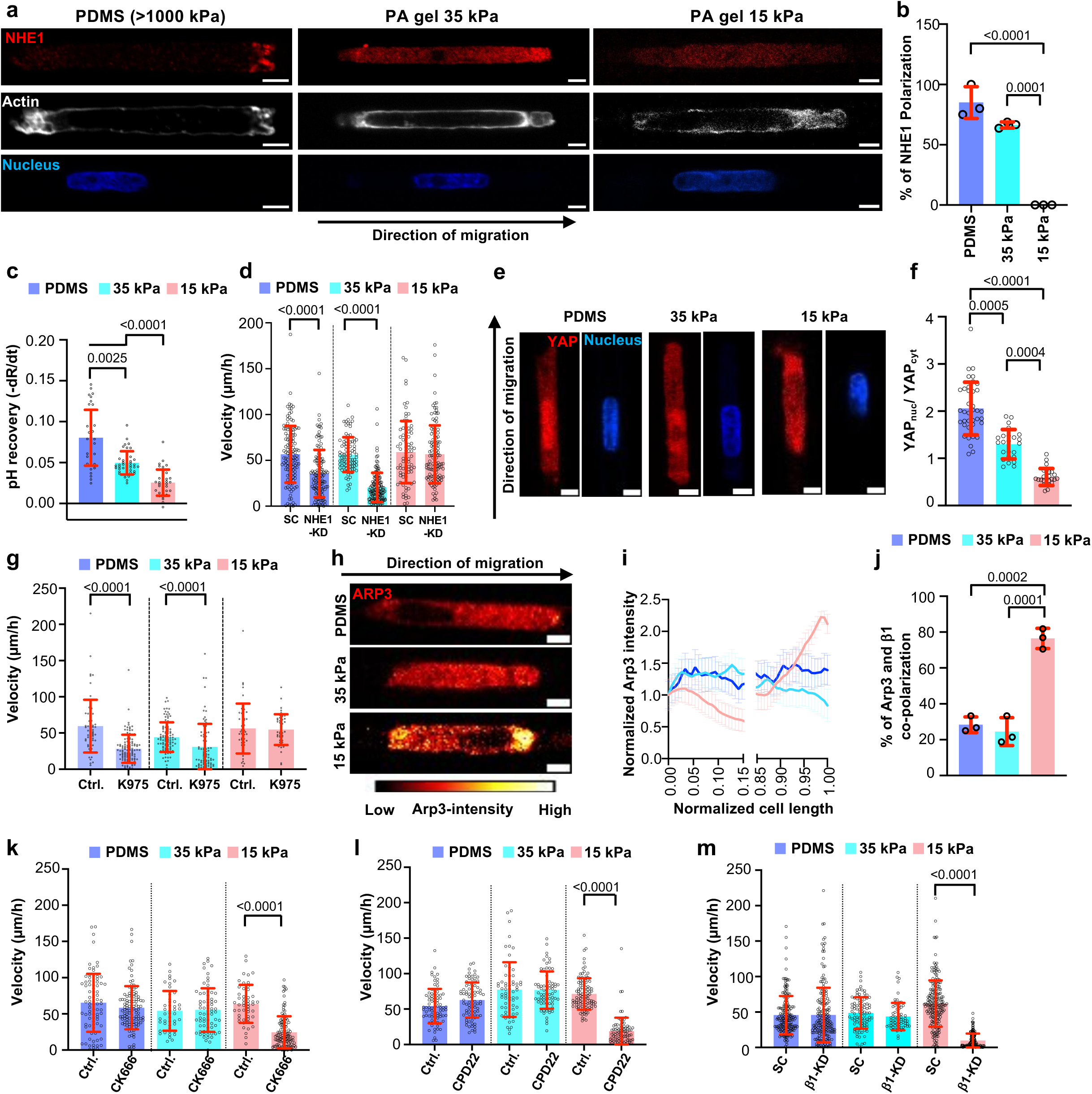
The roles of OEM and the cytoskeleton in cell motility within stiff and soft microchannels. **a**, Immunofluorescence images of cells in PDMS- or PA-based collagen I-coated microchannels of prescribed stiffness (35 or 15 kPa) stained for NHE1 (red), actin (white), and DNA (blue). Scale bars: 5 µm. **b**, Percentage of cells in PDMS- or PA-based collagen I-coated microchannels of prescribed stiffness displaying NHE1 polarization. Data represent mean±S.D. for n≥30 from 3 independent experiments. **c**, Measurements of NHE1 activity of cells on 2D collagen I-coated glass or PA gels of prescribed stiffness using NH_4_Cl prepulse technique by quantifying the pH_i_ recovery of pHrodo^TM^-loaded cells. Data represent mean±S.D. for n≥36 cells from 3 independent experiments. **d**, Migration velocity of SC and NHE1-KD cells in PDMS- or PA-based collagen I-coated microchannels. Data represent mean±S.D. for n≥82 cells from 3 independent experiments. **e**,**f**, Immunofluorescence images (e) and quantification (f) of nuclear-to-cytosolic YAP1 ratio of cells in PDMS- or PA-based microchannels of prescribed stiffness. Data represent mean±S.D. for n≥25 cells from 3 independent experiments. Scale bars: 5 µm. **g**, Cell migration velocity in PDMS- or PA-based collagen I-coated microchannels of prescribed stiffness in the presence of YAP1 inhibitor K975 or vehicle control (Ctrl.; DMSO). Data represent mean±S.D. for n≥64 cells from 3 independent experiments. **h**, Immunofluorescence images of cells in PDMS- or PA-based collagen I-coated microchannels stained for Arp3. Scale bars: 5 µm. **i**, ARP3 intensity normalized to the cell rear and plotted along the normalized cell length in confined cells. Data are the moving average±S.D. for n≥33 cells from 3 experiments. The *x-*axis is discontinued between 0.15 and 0.85 to highlight differences at the cell edges. **j**, Percentage of cells in PDMS- or PA-based collagen I-coated microchannels displaying Arp3 and β1-integrin front polarization. Data represent mean±S.D. for n≥30 from 3 independent experiments. **k-m**, Migration velocity of wild-type (k,l) or SC and β1-integrin knockdown (m) cells in PDMS- or PA-based collagen I-coated microchannels in the presence of Arp2/3 inhibitor CK666 (k) or ILK inhibitor CPD22 (l) or vehicle control (Ctrl.; DMSO). Data represent mean±S.D. for n≥46 cells from 3 independent experiments. Statistical significance was assessed by one-way ANOVA followed by Tukey’s (b,j), Kruskal-Wallis (c), Mann-Whitney (d, g, k-m), or unpaired t-test (f). Cell model: MDA-MB-231.

In view of these findings, we next examined whether migration in soft microenvironments is supported by cytoskeletal components rather than OEM. Arp3, a subunit of the seven-member actin-related protein 2/3 (Arp2/3) complex^18^, is strongly enriched at the cell leading edge in 15 kPa microchannels but appears diffused throughout the cell body in stiffer microchannels for MDA-MB-231, HOS, and triple negative breast cancer PDX cells (**Fig. 1h,i** and **Extended Data Fig. 1m-p**). Because actin-driven protrusion must be coordinated with integrin-based adhesions, which mechanically couple the extracellular matrix to the actin cytoskeleton during directional migration^21^, we next examined whether β1-integrin spatially colocalizes with Arp3 in soft microchannels. Indeed, Arp3 and β1-integrin co-polarize at the cell leading edge in softer, but not stiffer, microchannels (**Fig. 1j** and **Extended Data Fig. 1q**). Next, we examined integrin-linked kinase (ILK), an integrin-associated scaffolding protein that interacts with β1-integrin and links it to actin cytoskeleton^22^, thereby supporting cell migration. In line with this notion, β1-integrin, either alone or alongside ILK, preferentially polarizes at the cell front under all stiffness conditions (**Extended Data Fig. 1r,s**). Inhibition of Arp2/3 via CK666^18^ disrupts the front polarization of both β1-integrin and ILK in soft, but not stiff microchannels (**Extended Data Fig. 1t-v**). Conversely, β1-integrin knockdown^18^ abolishes Arp3 polarization (**Extended Data Fig. 1w**). These data illustrate that the functional loss of any of these cytoskeletal components disrupts the polarized localization of the others on soft substrates, highlighting a reciprocal dependency that reinforces front polarity. This spatial coordination is critical for sustained directional migration in softer microchannels, as inhibition of Arp2/3 with CK665, or the ILK-targeting compound CPD22^23^, as well as β1-integrin knockdown result in migration stalling (**Fig. 1k-m**). In contrast, cells in stiffer microchannels can bypass this requirement (**Fig. 1k-m**) in line with their reliance on the OEM^7,8^. Together, these findings demonstrate that substrate stiffness switches the cells migration mechanism from an actin cytoskeleton-dependent program in compliant microchannels to OEM-based migration in stiff microchannels. To further explore this, we set out to delineate the molecular pathway driving cancer cell migration in soft microenvironments.

## Cells use distinct migratory programs in soft versus stiff microchannels

MDA-MB-231 cells migrating in soft microchannels exhibit a higher membrane tension than cells on 2D substrates, as evidenced by their increased Flipper-TR lifetimes (**Extended Data Fig. 2a**). In line with the leading-edge polarization of Arp2/3, β1-integrin and ILK, and the established role of Arp2/3-dependent actin remodeling in regulating membrane tension^18,24^, increased membrane tension is detected only at the cell front of migrating cells in soft microchannels, and abolished by ILK inhibition via CPD22 (**Fig. 2a,b** and **Extended Data Fig. 2a**). As a control, we assessed membrane tension on 2D stiff (glass) and soft (15 kPa) substrates, and confirmed that hypotonic shock increases membrane tension under both conditions (**Extended Data Fig. 2b**). Changes in cell membrane tension lead to mechano/osmo-sensitive ion channel activation^18,25^, which in turn mediate calcium influx. In line with this notion, cells migrating in soft, but not stiff, confining microchannels display a markedly elevated calcium activity compared to cells in the adjacent 2D seeding region, as assessed by GCaMP6s imaging (**Fig. 2c,d** and **Supplementary Video 1,2**). In accord with this finding, treatment with the calcium chelator BAPTA-AM impairs the migration of MDA-MB-231 and HOS only in soft microchannels (**Fig. 2e** and **Extended Data Fig. 2c**). Given the central role of calcium in regulating confined migration in compliant microchannels, we aimed to identify the mechanosensitive ion channel(s) mediating the increased calcium activity. Pharmacological inhibition of TRPV4 via GSK2193874^18^ or inhibition of GsMTx4-senstive mechanosensitive ion channels, including Piezo1/2^26^, had no effect on confined migration (**Extended Data Fig. 2d**). In contrast, use of 2-APB, which inhibits the function of TRPC1, TRPM7 and inositol 1,4,5-triphosphate receptor^27^, markedly suppresses migration in soft, confining microchannels (**Extended Data Fig. 2d**). Moreover, specific inhibition of TRPM7 via fingolimod (FTY720)^27,28^ or TRPM7 knockout^27,28^ yields similar inhibitory effects for both MDA-MB-231 and SUM159 breast cancer cells (**Fig. 2f** and **Extended Data Fig. 2d,e**), revealing the critical role of TRPM7 in this process. In concert with this finding, pharmacological inhibition of TRPM7 nearly abolishes calcium activity inside soft microchannels, as evidenced by the markedly decreased GCaMP6s calcium spike frequency (spikes/min) (**Fig. 2g**). Moreover, inhibition of Arp2/3 via CK666, or ILK via CPD22 almost suppressed GCaMP6s spike frequency, suggesting that both ILK and Arp2/3 function as upstream regulators of membrane tension-induced TRPM7 activation and the consequent calcium influx (**Extended Data Fig. 2f**).

**Figure 2.**
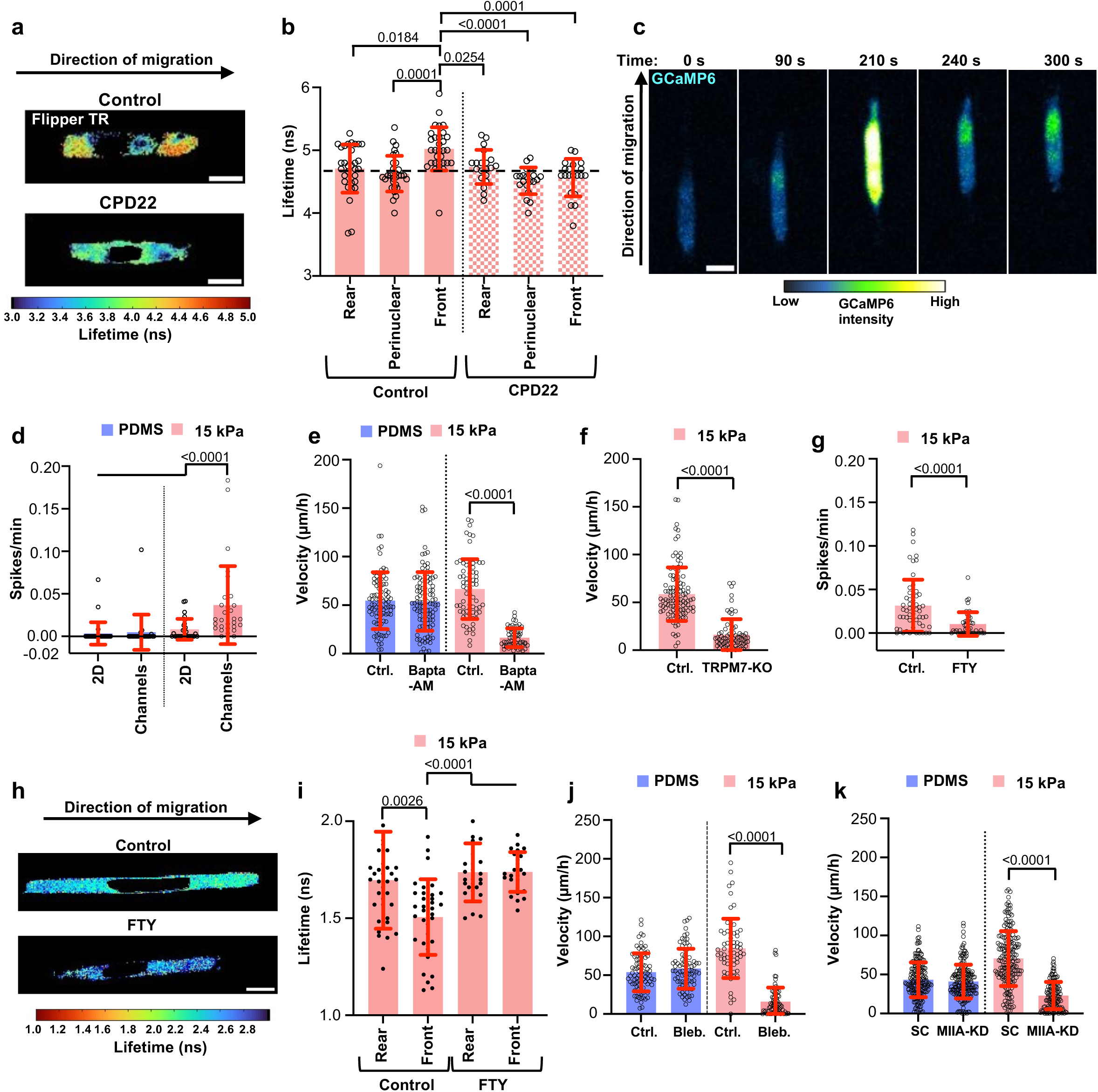
Cells use distinct migratory programs in soft versus stiff microchannels. **a**,**b**, Flipper-TR lifetime representative images (a) and quantification of membrane tension at the rear, perinuclear, and front (b) of cells in 15 kPa PA-based collagen I-coated microchannels in the presence of ILK inhibitor CPD22 or vehicle control (DMSO). Data are the mean±S.D. for n≥20 cells from 3 experiments. Scale bars: 5 µm. **c**, Representative images of GCaMP6 spikes over time for cells in 15 kPa PA-based collagen I-coated microchannels. Scale bar: 5 µm. **d**, GCaMP6 spikes/min of cells in PDMS- or 15 kPa PA-based collagen I-coated 2D area and microchannels. Data are mean±S.D. for n≥24 cells from 3 experiments. **e**, Cell migration velocity in PDMS- or 15 kPa PA-based collagen I-coated microchannels in the presence of the calcium chelator BAPTA-AM or vehicle control (Ctrl.; DMSO). Data represent mean±S.D. for n≥60 cells from 3 independent experiments. **f**, Migration velocity of CRISPR control or TRPM7-KO cells in 15 kPa PA-based collagen I-coated microchannels. Data represent mean±S.D. for n≥74 cells from 3 independent experiments. **g**, GCaMP6 spikes/min of cells in 15 kPa PA-based collagen I-coated microchannels in the presence of TRPM7 inhibitor FTY72 or vehicle control (Ctrl.; DMSO). Data are mean±S.D. for n≥45 cells from 3 experiments. **h**,**i**, Representative images (h) and quantification (i) of the lifetime of RhoA activity biosensor of the front and rear of RhoA2G-cells in 15 kPa PA-based collagen I-coated microchannels in the presence of TRPM7 inhibitor FTY720 or vehicle control (DMSO).. Scale bar: 5 µm. Data are mean±S.D. for n≥24 cells from 3 experiments. **j**, Cell migration velocity in PDMS- or 15 kPa PA-based collagen I-coated microchannels in the presence of blebbistatin or vehicle control (Ctrl.; DMSO). Data represent mean±S.D. for n≥51 cells from 3 independent experiments. **k**, Migration velocity of SC or MIIA-KD cells in PDMS- or 15 kPa PA-based collagen I-coated microchannels. Data represent mean±S.D. for n≥132 cells from 3 independent experiments. Statistical significance was assessed by one-way ANOVA followed by Tukey’s (b), Kruskal-Wallis (d,i), unpaired t-test (e), or Mann-Whitney (f,g,j,k). Cell model: MDA-MB-231.

In light of the association of TRPM7 with the regulation of RhoA-ROCK-myosin II pathway^27,28,29^, we first examined the potential contribution of RhoA to this process using confocal-fluorescence lifetime imaging microscopy (FLIM) coupled with a Förster resonance energy transfer (FRET)-based RhoA2G activity biosensor^18,30^. Cells inside soft, confining microchannels display increased RhoA activity relative to cells on 2D regions, as evidenced by the decreased donor fluorescence lifetimes, which is abrogated by TRPM7 inhibition (**Extended Data Fig. 2g**). Consistent with Arp2/3, β1-integrin and ILK leading-edge polarization, increased RhoA activity is detected at the anterior of migrating cells in soft microchannels, and abolished by TRPM7 inhibition via FTY (**Fig. 2h,i**). Moreover, inhibition of RhoA activity via a small-molecule Rho inhibitor or cell contractility via blebbistatin (20 µM) blocks migration in soft but not stiff microchannels (**Fig. 2j** and **Extended Data Fig. 2h**). Of note, partial inhibition of cell contractility using a low dose (5 µM) of blebbistatin does not alter migration in soft microchannels (**Extended Data Fig. 2i**). Interestingly, this partial inhibition increases migration velocities in stiff PDMS-based microchannels to match those measured in control cells within soft microchannels (**Extended Data Fig. 2i**), thereby enabling cells to perceive mechanically rigid substrates as compliant, consistent with the reduced contractility on softer matrices^31,32^. Myosin IIA (MIIA), but not MIIB, knockdown^27^ suppresses migration in soft, but not stiff, microchannels (**Fig. 2k** and **Extended Data Fig. 2j**). Moreover, dual knockdown of MIIA and MIIB isoforms does not further decrease motility relative to MIIA knockdown alone (**Extended Data Fig. 2k**), supporting MIIA as the primary driver of contractility-dependent migration under compliant confinement. In contrast to the dominant role of the OEM in stiff microchannels, its key regulator NHE1 is neither polarized at the cell front nor functionally supporting migration in soft microchannels. Together, these results establish an OEM-independent molecular program supporting migration in straight, confining, soft microchannels, in which Arp2/3-ILK-β1-integrin coupling increases membrane tension, which in turn promotes TRPM7-mediated calcium influx, leading to RhoA activation and MIIA-dependent contractility.

## Distinct metabolic requirements for cell motility in soft versus stiff microchannels

Given the mechanistically distinct migratory programs adopted by cancer cells in soft versus stiff microchannels, we next examined whether these divergent mechanisms are accompanied by shifts in cell’s bioenergetic requirements. To this end, we inhibited the two major bioenergetic pathways supporting cellular ATP production, glycolysis and mitochondrial OxPHOS. Glycolysis inhibition via 2-deoxy-D-glucose (2-DG) suppresses cell migration in stiff, but not soft microchannels of 15 kPa (**Fig. 3a**). Interestingly, inhibition of OxPHOS using oligomycin exerts the exact opposite effect by selectively impairing migration in soft, but not stiff microchannels, as observed for both MDA-MB-231 breast cancer and HOS cells (**Fig. 3a** and **Extended Data Fig. 3a**). The inhibitory effects of OxPHOS on confined migration inside soft microchannels are attributed to the disruption of Arp3 and β1-integrin polarization at the cell leading edge (**Fig. 3b**). Of note, this polarization pattern remains unaffected by glycolysis inhibition (**Fig. 3b**). Similar observations were made for even softer (7 kPa) polyacrylamide-based microchannels (**Extended Data Fig. 3b-d**). In distinct contrast, glycolysis, but not OxPHOS, inhibition impairs OEM-dependent migration in stiff PDMS-based microchannels by interfering with NHE1 polarization at the cell anterior (**Fig. 3c,d**). We further demonstrated the dependence of migration on glycolysis, but not OxPHOS, for cells inside PA-based microchannels with a stiffness of 35 kPa (**Extended Data Fig. 3e**).

**Figure 3.**
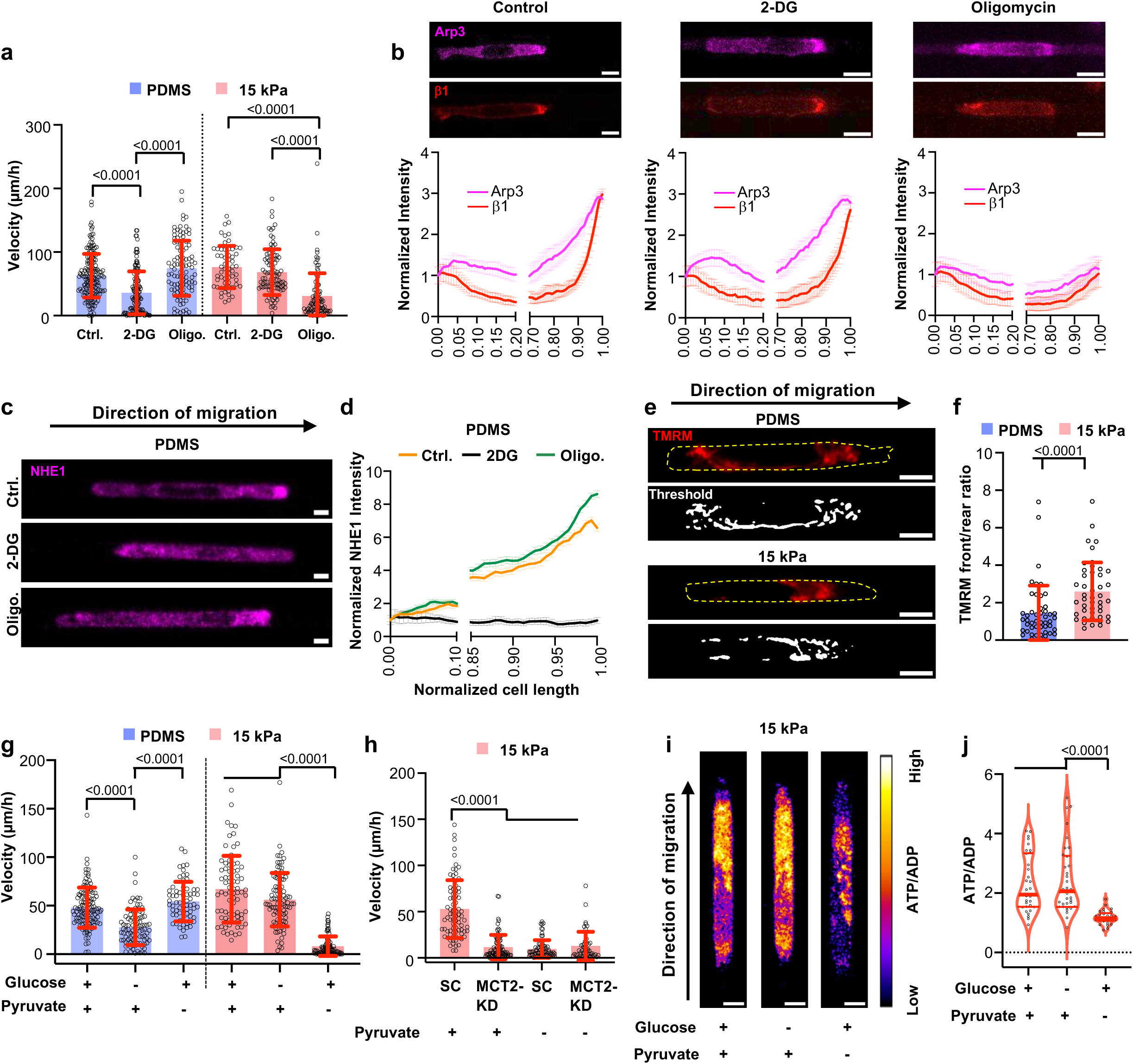
Distinct metabolic requirements for cell motility in soft versus stiff microchannels. **a**, Cell migration velocity in PDMS- or 15 kPa PA-based collagen I-coated microchannels in the presence of 2-DG or oligomycin (Oligo.) or vehicle control (Ctrl.; DMSO). Data represent mean±S.D. for n≥62 cells from 3 independent experiments. **b**, Immunofluorescence images, and intensity normalized to the cell rear and plotted along the normalized cell length of cells in 15 kPa PA-based collagen I-coated microchannels stained for Arp3 (magenta) and β1-integrin (red) in the presence of 2-DG or oligomycin or vehicle control (DMSO). Scale bars: 5 µm. Data are the moving average±S.D. for n≥25 cells from 3 experiments. The *x-* axis is discontinued between 0.20 and 0.70 to highlight differences at the cell edges. **c**,**d** Immunofluorescence images (c), and NHE1 intensity normalized to the cell rear and plotted along normalized cell length (d) of cells in PDMS-based collagen I-coated microchannels stained for NHE1 (magenta) in the presence of 2-DG or oligomycin (Oligo.) or vehicle control (Ctrl.; DMSO). Data are the moving average±S.D. for n≥33 cells from 3 experiments. The *x-* axis is discontinued between 0.10 and 0.85 to highlight differences at the cell edges. Scale bars: 5 µm. **e**,**f**, Representative immunofluorescence and thresholded images (e) and front/rear ratio (f) of TMRM (red) intensity in cells migrating in PDMS-based or 15 kPa PA-based collagen I-coated microchannels. Data represent mean±S.D. for n≥48 cells from 3 independent experiments. Scale bars: 5 µm. **g**, Cell migration velocity in PDMS-or 15 kPa PA-based collagen I-coated microchannels in medium with or without glucose or pyruvate. Data represent mean±S.D. for n≥66 cells from 3 independent experiments. **h**, Migration velocity of SC or MCT2-KD cells in 15 kPa PA-based collagen I-coated microchannels in the presence of medium with or without pyruvate. Data represent mean±S.D. for n≥49 cells from 3 independent experiments. **i**,**j**, Representative immunofluorescence images (i) and quantification (j) of PercevalHR (ATP/ADP) ratio of cells co-expressing Perceval HR and pHRed inside 15 kPa microchannels in medium with or without glucose or pyruvate. Data represent mean±S.D. for n≥29 cells from 3 independent experiments. Red lines indicate the median (thick) and quartile (thin). Scale bars: 5 µm. Statistical significance was assessed by Kruskal-Wallis (a,g,h), unpaired t-test (f), or one-way ANOVA followed by Tukey’s (l). Cell model: MDA-MB-231.

To further establish the critical role of mitochondrial activity in confined migration in soft microchannels, we depleted mitochondria by treating Parkin-overexpressing cells with trifluoromethoxy carbonylcyanide phenylhydrazone (FCCP) for 3-6 h^33^, which induces mitochondrial damage and their subsequent clearance by Parkin (**Extended Data Fig. 3f,g**). This intervention abrogates confined migration in soft microchannels as well as Arp3 polarization, consistent with the role of OxPHOS in this process (**Extended Data Fig. 3h-j** and **Supplementary Video 3)**. Remarkably, despite mitochondrial depletion, cells retain their migratory capacity in stiff microchannels, demonstrating that OEM-driven migration can operate independently of mitochondria themselves (**Extended Data 3h**, **Supplementary Video 4**). In line with the dispensable role of mitochondria in cell motility inside stiff microchannels, mitochondrial depletion fails to alter NHE1 leading-edge polarization (**Extended Data Fig. 3k**).

The involvement of OxPHOS and mitochondrial function in supporting migration in soft microchannels was further substantiated by Tetramethylrhodamine, methyl ester (TMRM) staining, which reveals that mitochondria are polarized toward the cell leading edge in soft, but not stiff, channels (**Fig. 3e,f** and **Supplementary Video 5**). Importantly, TMRM polarization at the cell anterior coincides with the direction of migration. As such, upon cell reversal, TMRM spatial localization shifts to the cell rear (new leading edge), whereas mitochondria are equally distributed to the cell poles in stalling cells (**Extended Data Fig. 3l,m**). Consistent with previous work on 2D surfaces^34^, we observed stiffness-dependent differences in mitochondrial morphology, with more elongated mitochondria on stiffer substrates (**Extended Data Fig. n,p**). In contrast, no difference in mitochondrial form factor was detected for cells inside stiff versus soft microchannels (**Extended Data Fig. o,p**). These findings suggest that, during confined migration in soft microchannels, mitochondrial spatial organization and activity, rather than morphology, may play a more prominent functional role.

Active mitochondria have been reported to co-polarize with actin at the leading edge of migrating cells^35^, while ILK knockdown in MDA-MB-231 disrupts mitochondrial localization^36^, supporting close coordination between mitochondrial positioning and actin-adhesion dynamics^35,36^. Consistent with these observations, live imaging of cells expressing subunit 9 of mitochondrial ATPase (SU9)-GFP-expressing cells, mitochondria consistently localize just posterior to Arp3 and β1-integrin at the leading edge of migrating cells in soft microchannels (**Extended Data Fig. 3q,r**). These data suggest that mitochondrial positioning may support the establishment or maintenance of Arp3 and β1-integrin during migration in soft microchannels. In concert with this notion, inhibition of Arp2/3 via CK666, calcium chelation via BAPTA-AM, or TRPM7 activity via FTY720, disrupt mitochondrial front polarization in soft microchannels (**Extended Data Fig. 3s-v**). Importantly, these data point to a potential feedback loop in which mitochondrial leading-edge localization is required for Arp2/3 polarization and vice versa. This reciprocal interaction between mitochondria and Arp2/3 is further sustained by TRPM7-mediated calcium influx, thereby facilitating migration in soft microenvironments. Together, these findings establish a mechano-metabolic link that depends on substrate stiffness: an OEM-based, glycolysis-dependent mechanism predominates in stiff microchannels, while a mitochondria-dependent, cytoskeletal- and TRPM7-driven program supports migration in soft confining spaces.

## Monocarboxylate Transporter 2 (MCT2)-driven pyruvate uptake fuels mitochondria-based migration in soft microchannels

To delineate the specific metabolic substrate that fuels mitochondrial function during confined migration in soft versus stiff microenvironments, we selectively inhibited the three principal inputs into the tricarboxylic acid (TCA) cycle. Fatty-acid oxidation was suppressed via etomoxir, glutaminolysis was blocked using the glutaminase inhibitor CB-839, and mitochondrial pyruvate import was disrupted using the mitochondrial pyruvate carrier (MPC) inhibitor UK-5099. In stiff PDMS microchannels, pharmacologic inhibition of these pathways did not alter cell migratory capacity (**Extended Data Fig. 4a**), consistent with the dispensable role of mitochondria in this process. In contrast, in soft microchannels, only UK-5099 impaired confined migration (**Extended Data Fig. 4b**), indicating that pyruvate-driven OxPHOS is the primary metabolic driver of migration in soft microenvironments. Along these lines, removal of pyruvate selectively abrogated migration in soft microchannels **(Fig. 3g)** and was accompanied by a marked decrease in TMRM intensity, consistent with reduced mitochondrial activity (**Extended Data Fig. 4c**). On the other hand, and in concert with the glycolytic dependence of OEM-driven migration within stiff confining spaces, complete removal of glucose from the medium reduced migration in stiff, but not soft, microchannels (**Fig. 3g**). These results demonstrate that cells migrating in soft confining spaces depend on pyruvate-driven mitochondrial metabolism but not glycolysis, indicating a functional uncoupling of glycolysis from OxPHOS.

In view of the role of MCT2 as the membrane transporter responsible for pyruvate uptake^37^, we knocked down MCT2 (**Extended Data Fig. 4d,e**) and found that this molecular intervention markedly impaired migration in soft, but not stiff, microchannels, reducing velocities to levels comparable to those observed in pyruvate-free media (**Fig. 3h** and **Extended Data Fig. 4f**). This reduction in migration velocity is corroborated by findings showing that MCT2 knockdown impaired Arp3 and β1-integrin front co-polarization in soft microchannels (**Extended Data Fig. 4g,h**), disrupting the cytoskeletal organization required for efficient migration under soft confinement. Together, these findings indicate that pyruvate-dependent mitochondrial function is required to establish the front Arp3/β1-integrin polarity needed for migration in soft microchannels. Depletion of pyruvate, but not glucose, decreased the ATP/ADP ratio in soft microchannels as measured with PercevalHR, further supporting pyruvate-fueled OxPHOS as the predominant energetic source sustaining migration under this condition (**Fig. 3i,j**). Because PercevalHR is pH-sensitive, we used cells co-expressing PercevalHR and pHRed to confirm that intracellular pH remains unchanged across conditions (**Extended Data Fig. 4i**).

## NHE1 overexpression and activation promotes OEM-based migration in soft microchannels

In light of the absence of NHE1 polarization and reduced activity in soft substrates, we next asked whether restoring NHE1 function could promote OEM-based migration in soft microenvironments. To this end, NHE1 overexpression^18^ enhances migration velocity only in soft microchannels (**Fig. 4a** and **Extended Data Fig. 5a**). In distinct contrast to wild-type cells, treatment of NHE1-overexpressing cells with inhibitors of Arp2/3 (CK666) or ILK (CPD22) or contractility (blebbistatin) does not reduce migration in soft microchannels (**Fig. 4b**), mimicking the behavior of wild-type cells albeit in stiff confining spaces. Interestingly, NHE1-overexpressing MDA-MB-231 cells are able to migrate in soft microchannels and inside 3D collagen gels even after complete disruption of actin polymerization via latrunculin A (LatA) (**Fig. 4c,d**, **Extended Data Fig. 5b** and **Supplementary Video 6-8**), revealing a mechanistic switch to OEM-based migration. This is further corroborated by findings showing that about 60% of untreated NHE1-overexpressing cells exhibit leading-edge polarization of NHE1 as opposed to the absence of any NHE1 enrichment at the anterior of wild-type control cells in soft microchannels (**Extended Data Fig. 5c,d**). The switch to OEM-based migration was further validated using SUM159 breast cancer and HOS cells (**Extended Data Fig. 5e-h**), as evidenced by their ability to migrate in soft microchannels even after complete disruption of actin polymerization.

**Figure 4.**
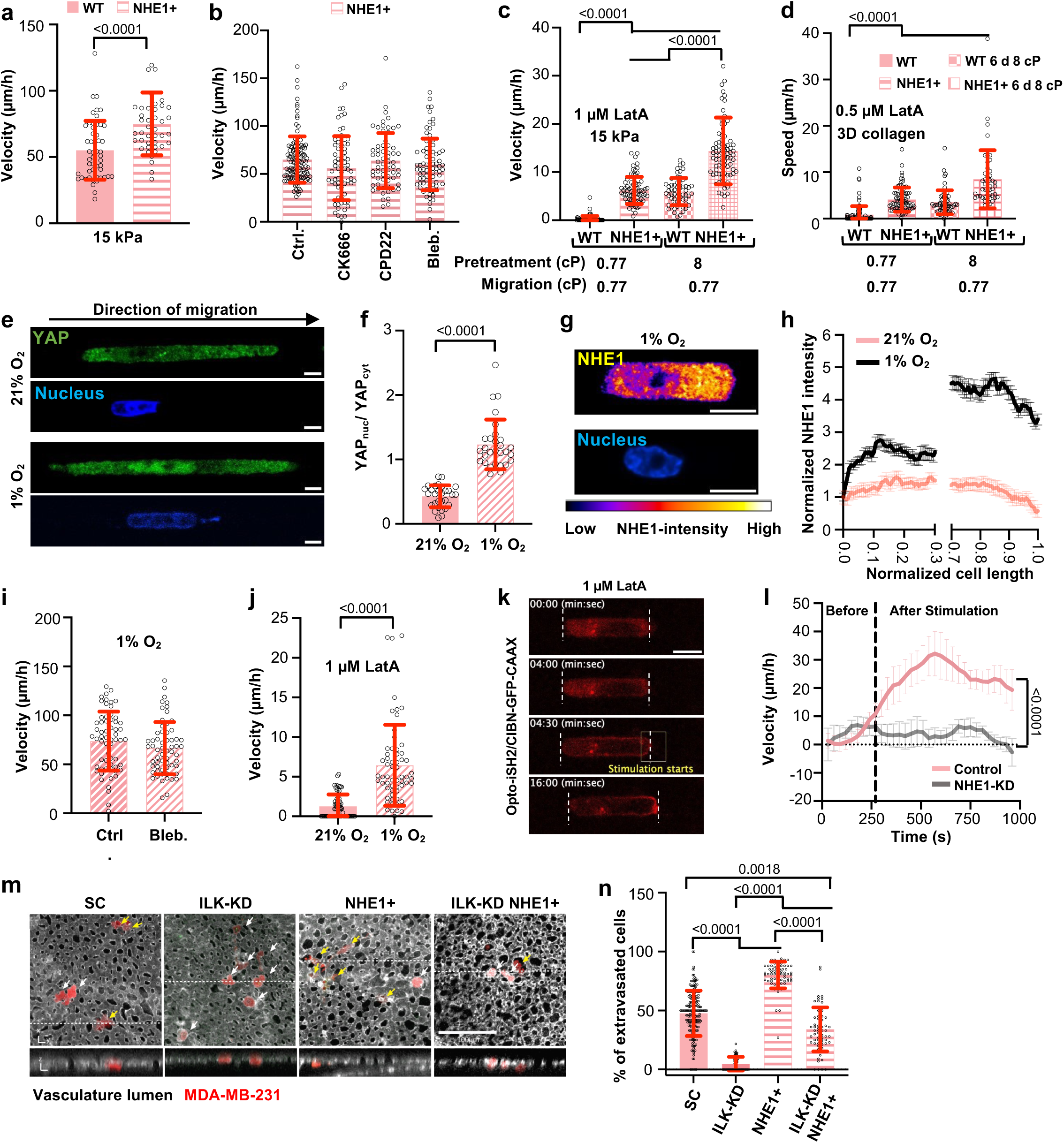
NHE1 overexpression and activation promotes OEM-based migration in soft microchannels. **a**, Migration velocity of wild-type and NHE1-overexpressing cells in 15 kPa PA-based collagen I-coated microchannels. Data are mean±S.D. for n≥42 cells from 3 independent experiments. **b**, Migration velocity of NHE1-overexpressing cells in 15 kPa PA-based collagen I-coated microchannels in the presence of CK666, CPD22, blebbistatin, or vehicle control (Ctrl.; DMSO). Data are mean±S.D. for n≥68 cells from 3 independent experiments. **c**,**d**, Migration velocity (c) or speed (d) of wild-type and NHE1-overexpressing cells pretreated for 6 days in 0.77 or 8 cP media, seeded in 15 kPa PA-based collagen I-coated microchannels (c) or in 3D collagen (d), and treated with LatA during migration. Data are mean±S.D. for n≥49 cells from 3 independent experiments. **e**,**f**, Immunofluorescence images (e) and quantification (f) of nuclear-to-cytosolic YAP1 ratio of cells preconditioned to normoxia or hypoxia for 48 h and in 15 kPa PA-based microchannels. Data represent mean±S.D. for n=29 cells from 3 independent experiments. Scale bars: 5 µm. **g**,**h**, Representative immunofluorescence image (g) and NHE1 intensity normalized to the cell rear and plotted along the normalized cell length (h) for cells pretreated in hypoxia for 48 h. Data are the moving average ± S.D. for n≥26 cells from 3 experiments. The *x-*axis is discontinued between 0.3 and 0.7 to highlight differences at the cell edges. Scale bar: 5 µm. **i,j,** Cell migration velocity after exposure to hypoxia (i) or to normoxia or hypoxia (j) for 48 h in 15 kPa PA-based collagen I-coated microchannels in the presence of blebbistatin (i) or LatA (j) or vehicle control (Ctrl.; DMSO). Data are the mean ± S.D. for n≥60 cells from 3 experiments. **k**, Time-lapse montage of a representative cell expressing opto-iSH2 and CIBN-GFP-CAAX and migrating in 15 kPa PA-based collagen I-coated microchannels before and after light stimulation at the cell leading edge in a region enclosed by the yellow box. Dashed white lines indicate the cell edges. Scale bar: 5 µm. **l**, Migration velocity over time of control and NHE1-KD opto-iSH2 CIBN-GFP-CAAX cells in 15 kPa PA-based collagen I-coated microchannels before and after light stimulation. Data are mean±S.E.M. for n=11 (control) or n=5 (NHE1-KD) cells from 3 independent experiments. **m,** Representative images showing imaging fields (XY view in top panels or ZY view in lower panels) of chicken CAM vasculature with extravasating SC, NHE1-overexpressing, ILK-KD, and ILK-KD NHE1-overexpressing mCherry cells. Yellow arrows point to the extravasated cells, white arrows point to the cells that are still inside the vasculature lumen. Scale bar: 100 µm. **n,** Percentage (%) of extravasated SC, NHE1-overexpressing, ILK-KD, and ILK-KD NHE1-overexpressing mCherry cells in the chick embryo model shown in m. Data are mean±S.D. of an independent imaging field for n≥9 animals from 3 independent experiments. Statistical significance was assessed by unpaired t-test (f) after lognormal transformation (a), Kruskal-Wallis (c,d,n), or Mann-Whitney (j). Cell model: MBA-MB-231

Cancer cells, in their journey to invade and metastasize, must adapt and react to different mechanochemical cues within the tumor microenvironment. We hypothesized that among these stimuli, those promoting NHE1 expression and/or activity, such as hypoxia^38^ and elevated viscosity^18^, may trigger the switch from actomyosin cytoskeleton-based to OEM-dependent cell migration in soft substrates. To test this hypothesis, we first examined the effect of hypoxia on YAP nuclear localization, and NHE1 polarization and activity, as well as cell motility in soft microchannels. Cells preconditioned to hypoxia (1% O₂ for 48 h), as opposed to normoxia, displayed higher YAP nuclear localization in soft microchannels, which was accompanied by increased NHE1 polarization at the leading edge and elevated NHE1 activity (**Fig. 4e-h** and **Extended Data Fig. 5i**). Moreover, hypoxia promoted contractility-independent migration in soft microchannels (**Fig. 4i)**. Importantly, migration of hypoxia-preconditioned cells was detected even after complete disruption of actin polymerization via LatA under soft confinement (**Fig. 4j**), supporting a hypoxia-mediated switch to OEM-based migration.

In agreement with work showing that elevated extracellular fluid viscosity (8 cP) enhances NHE1 activity^18,24^, and cancer cell migration^18^, wild-type cancer cell preconditioning to 8 cP for 6 days increased NHE1 activity (**Extended Data Fig. 5j**) and promoted faster migration in soft microchannels (**Extended Data Fig. 5k**). It is noteworthy that viscosity-preconditioned wild-type cells are capable of migrating in soft microchannels and 3D collagen gels even in the absence of actin polymerization following LatA treatment, reaching migration velocities and speeds comparable to those of naive NHE1-overexpressing cells (**Fig. 4c,d**, **Extended Data Fig. 5b** and **Supplementary Video 9**). Remarkably, preconditioning of NHE1-overexpressing cells to elevated viscosity for 6 days further increased confined migration in soft microchannels under LatA treatment, suggesting an additive effect between upregulation of NHE1 expression and activity (**Fig. 4c,d** and **Supplementary Video 10,11**). In sum, these data represent the first demonstration of OEM-based migration in soft microchannels and 3D collagen gels.

In view of our study^39^ revealing a direct interaction and a bidirectional regulatory loop between NHE1 and phosphoAkt, we developed an optogenetic tool to control the spatiotemporal activation of Akt. Using the cryptochrome (Cry2)/CIBN light-gated dimerization system^40^, we selectively activated Akt at the leading edge of migrating cells^39,41^. Optogenetic activation of Akt at the leading edge of wild-type MDA-MB-231 cells migrating in soft microchannels in the presence of LatA, rescues their migratory capacity, which is abolished by NHE1 knockdown (**Fig. 4k,l**). These findings indicate that localized Akt activation is sufficient to promote actin-independent, NHE1-dependent OEM-based migration under compliant confinement.

To further establish the critical involvement of OEM-based migration in soft microenvironments, we knocked down ILK in scramble control and NHE1-overexpressing MDA-MB-231 cells (**Extended Data Fig. 5l**). In accord with the role of ILK in regulating actin stress fiber formation and focal adhesion maturation^42,43^, ILK knockdown cells exhibit a complete loss of migratory capacity in 3D collagen gels, whereas NHE1 overexpression is sufficient to partially restore their motility (**Extended Data Fig. 5m**). These findings were validated *in vivo* using chick embryos, whose tissue stiffness varies between 0.5 and 1 kPa^44^. Under these compliant conditions, ILK knockdown cells fail to extravasate, while NHE1 overexpression partially rescues their extravasation ability (**Fig. 4m,n**). Of note, NHE1-overexpressing cells display a higher migratory and extravasation potentials *in vitro* and *in vivo*. Taken together, these data suggest that activating the OEM can compensate for impaired actin-integrin coupling in soft microenvironments.

## Cell intracellular rewiring overrides stiffness-dependent metabolic adaptation

Having established that induction of NHE1 polarization and activity restores OEM-based migration in soft microchannels, we next investigated how this cell intracellular rewiring affects its bioenergetic requirements. While wild-type cell migration in soft confining spaces relies on OxPHOS, overexpression of NHE1 via either molecular or chemical (hypoxia treatment for 48 h)^39^ means is sufficient to switch the cell metabolic requirements to glycolysis, as evidenced by a pronounced reduction in motility upon 2-DG treatment (**Fig. 5a** and **Extended Data Fig. 6a**). Thus, NHE1-overexpressing cells in soft microchannels mirror the metabolic profile of cells in stiff surfaces (**Fig. 3a**). In line with these findings, while mitochondria are front-polarized in wild-type cells migrating in 15 kPa microchannels, this polarization pattern is lost in NHE1-overexpressing cells (**Extended Data Fig. 6b,c**). This is accompanied by reduced mitochondrial activity in NHE1-overexpressing cells compared to wild-type cells migrating in 15 kPa microchannels, as indicated by lower TMRM intensity (**Extended Data Fig. 6d**). We next asked whether disrupting NHE1 function in stiff microchannels produces the reciprocal metabolic switch. While control cells primarily depend on glycolysis for migration within stiff microchannels (**Fig. 3a**), NHE1 knockdown suppresses cell motility and shifts their metabolic dependence to OxPHOS, as evidenced by their sensitivity to oligomycin (**Extended Data Fig. 6e**). Along these lines, reducing NHE1 front polarization and activity, via cell treatment with the YAP inhibitor K975, suppresses cancer cell migration in stiff microchannels, to an extent similar to that observed under glucose deprivation (**Extended Data Fig. 6f**). Of note, no additive effect is detected upon K975 treatment in glucose-deprived media (**Extended Data Fig. 6f**). Remarkably, while MCT2 knockdown does not alter migration in stiff microchannels, this molecular intervention coupled with either YAP inhibition or glucose deprivation nearly stalls motility (**Extended Data Fig. 6f**), thereby suggesting a shift to OxPHOS only when glucose supply is limited or NHE1 activity and polarization are disrupted. Together, these findings demonstrate that intracellular rewiring can override external stiffness cues and shift the metabolic requirements of migrating cells irrespective of the mechanical properties of their microenvironment.

**Figure 5.**
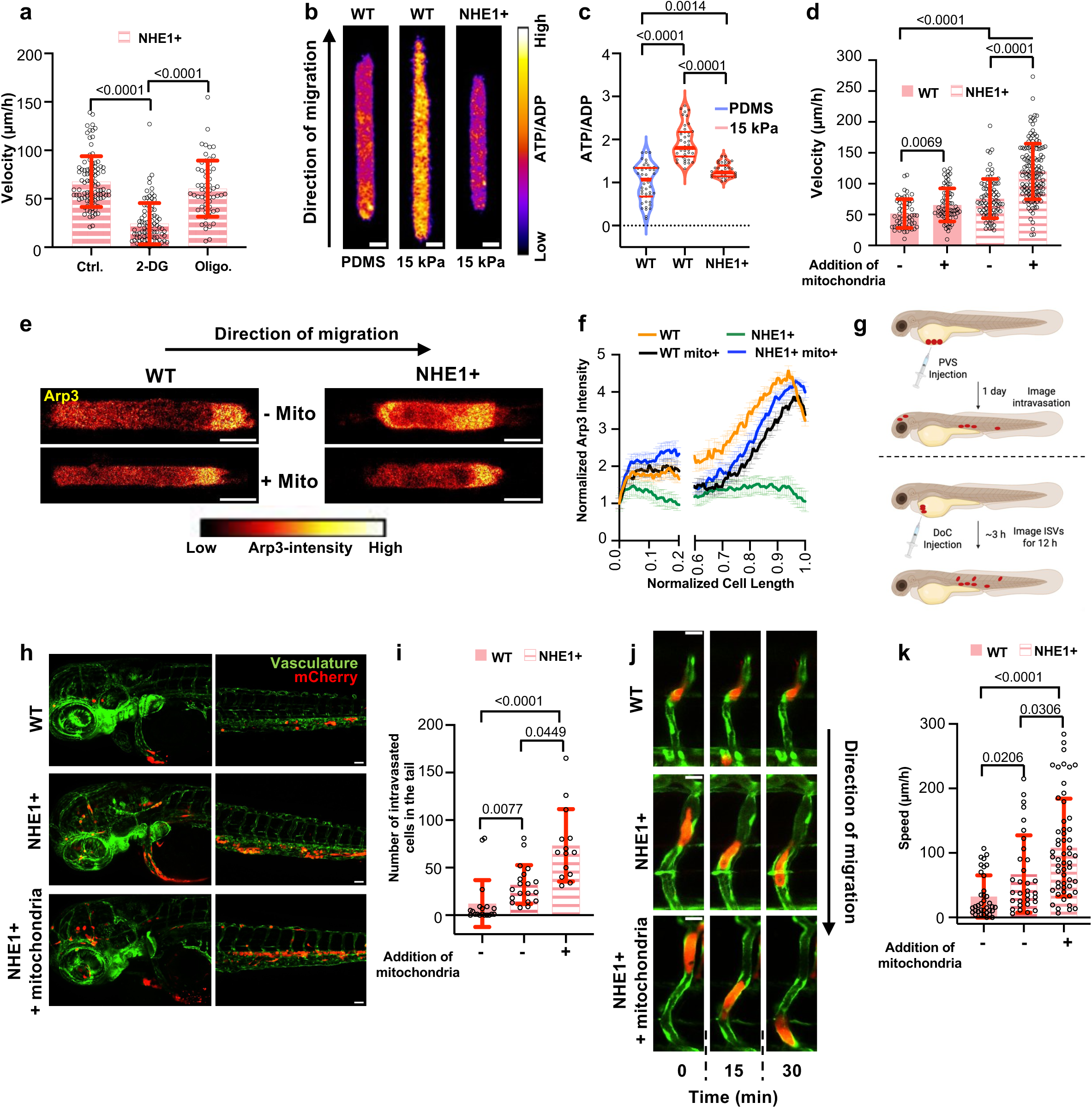
Cell intracellular rewiring overrides stiffness-dependent metabolic adaptation. **a**, Migration velocity of NHE1-overexpressing cells in 15 kPa PA-based collagen I-coated microchannels in the presence ce of 2-DG or oligomycin (Oligo.) or vehicle control (Ctrl.; DMSO). Data represent mean±S.D. for n≥58 cells from 3 independent experiments. **b**,**c**, Representative immunofluorescence images (b) and quantification (c) of PercevalHR (ATP/ADP) ratio of wild-type or NHE1-overexpressing cells co-expressing Perceval HR and pHred in PDMS- or 15 kPa PA-based microchannels. Data represent mean±S.D. for n≥39 cells from 3 independent experiments. Red lines indicate the median (thick) and quartile (thin). Scale bars: 5 µm. **d**, Migration velocity of wild-type and NHE1-overexpressing cells in 15 kPa PA-based collagen I-coated microchannels with or without exogenous mitochondrial addition. Data represent mean±S.D. for n≥59 cells from 3 independent experiments. **e**, Arp3 immunofluorescence images of wild-type and NHE1-overexpressing cells in 15 kPa PA-based collagen I-coated microchannels with or without exogenous mitochondrial addition. Scale bars: 5 µm. **f**, Arp3 intensity normalized to the cell rear and plotted along the normalized cell length in confined wild-type and NHE1-overexpressing cells with or without exogenous mitochondrial addition. Data are the moving average ± S.D. for n≥23 cells from 3 experiments. The *x-* axis is discontinued between 0.2 and 0.6 to highlight differences at the cell edges. **g**, Schematic showing the workflow of perivitelline space (PVS) and Duct of Cuvier (DoC) injections in the zebrafish to assess cell intravasation and migration inside intersegmental vessels (ISVs), respectively. **h**, Representative zebrafish vasculature (green) containing mCherry wild-type or NHE1-overexpressing with or without exogenous mitochondrial addition (red) 1 day after PVS injection. Scale bars: 50 µm. **i,** number of mCherry wild-type or NHE1-overexpressing cells in the presence or absence of exogenous mitochondrial addition intravasated to the tail in a zebrafish model 1 day after injection in the PVS. Data are mean±S.D. for n≥14 fish from 4 experiments. **j**,**k**, Representative zebrafish images (j) and migration speed (k) of mCherry-tagged wild-type or NHE1-overexpressing cells (red) with or without exogenous mitochondrial addition migrating in ISVs (green). Data are mean±S.D. for n≥33 cells from 3 independent clutches. Scale bars: 20 µm. Statistical significance was assessed by Kruskal-Wallis (a,c,i,k), one-way ANOVA followed by Tukey’s after log-normal transformation (d). Cell model: MDA-MB-231.

To further establish the presence of a bidirectional crosstalk between NHE1 and glycolysis, the NADH/NAD^+^ Peredox biosensor revealed that that pharmacological inhibition of NHE1 activity via EIPA, reduces the glycolysis-associated metabolic state in stiff, PDMS-based microchannels (**Extended Data Fig. 6g,h**). Conversely, NHE1 overexpression enhances glycolysis-associated NADH signature under soft confinement (**Extended Data Fig. 6g,h**), consistent with the role of NHE1 to bias cells towards a more glycolysis-supported energetic state even in compliant microenvironments. Importantly, PercevalHR ATP/ADP measurements revealed that migration in compliant (15 kPa) microchannels is more energy intensive than in stiffer (35 kPa) PA-based (**Extended Data Fig. 6i**) or PDMS-based ones (**Fig. 5b,c**). However, NHE1 overexpression compared to wild-type controls is associated with a decreased ATP/ADP ratio in soft microchannels (**Fig. 5b,c** and **Extended Data Fig. 6i**) despite increased cell velocities (**Extended Data Fig. 6j**). Consistent with these observations, seahorse analysis indicates that NHE1-overexpressing cells have a lower total ATP production rate than wild-type cells (**Extended Data Fig. 6k**). Because Ki67 staining shows no differences in proliferation between wild-type and NHE1-overexpressing cells, we rule out altered proliferative capacity as a driver of these metabolic changes (**Extended Data Fig. 6l**). Together, these findings reveal that activation of the OEM alone is sufficient to drive intracellular metabolic rewiring, enabling cells to migrate while operating at a lower energetic state. These findings suggest that distinct migration modes are associated with different energetic demands, with OEM engagement enabling cells to maintain, or even enhance, migratory capacity despite reduced ATP production.

The reduced mitochondrial activity in NHE1-overexpressing cells (**Extended Data Fig. 6d**) prompted us to investigate whether enhancing mitochondrial function would alter their migratory behavior. This point is particularly relevant as cancer cells *in vivo* do not operate in isolation but interact extensively with their surrounding microenvironment, including stromal cells such as cancer-associated fibroblasts (CAFs), which are known to transfer lipids, metabolites, and mitochondria to tumor cells to support their survival and invasion^45,46^. These metabolic exchanges are particularly critical under stress conditions such as matrix confinement, where tumor cells rely on stromal support to maintain bioenergetic and redox balance^45,46^. To this end, we isolated mitochondria from CAFs and introduced them into MDA-MB-231 cells. Successful mitochondrial incorporation was confirmed by increased MitoTracker intensity, elevated oxygen consumption rate (OCR), and higher mitochondrial DNA content in both wild-type and NHE1-overexpressing cells (**Extended Data Fig.7a-c**). Mitochondrial uptake was further validated by transplanting isolated mitochondria from mito-dsRED2-labeled CAFs into unlabeled recipient MDA-MB-231 cells, revealing dsRED2-positive mitochondria in the recipient cells (**Extended Data Fig. 7d**). While mitochondrial addition only modestly enhances motility in wild-type cells, it markedly increases migration velocity (**Fig. 5d**), restores mitochondrial front polarization and activity in NHE1-overexpressing cells in soft microchannels (**Extended Data Fig.7e-h**). In accord with the role of mitochondria in regulating Arp3 polarization (**Fig. 3b**), exogenous mitochondrial addition in NHE1-overexpressing cells also restores Arp3 front polarization in soft microchannels to the levels of wild-type cells (**Fig. 5e,f**). As a control, we confirmed that exogenous addition of mitochondria does not influence OEM-mediated motility, as migration velocities remain unchanged after actin disruption via LatA (**Extended Data Fig.7i**). Moreover, NHE1 polarization is unaltered in both wild-type and NHE1-overexpressing cells in soft microchannels irrespective of mitochondrial addition (**Extended Data Fig.7j**). To validate these findings, as an alternative to exogenous mitochondrial transplantation, we directly activated endogenous mitochondrial activity via cell treatment with an activator of the mitochondrial calcium uniporter (MCU), kaempferol^47^. This treatment was sufficient to enhance mitochondrial activity (**Extended Data Fig. 7k**), increase Arp3 front polarization (**Extended Data Fig. 7l,m**), and promote migration inside soft microchannels (**Extended Data Fig. 7n**), resulting in a hyper-motile cell phenotype. Using the zebrafish model, we further demonstrate that NHE1-overexpressing relative to wild-type cells, exhibit enhanced intravasation to the tail and brain after one day post-injection into the perivitelline space (**Fig. 5g-i** and **Extended Data Fig.7o**), as well as increased migration speed in the in the narrow intersegmental vessels (ISVs) (**Fig. 5g,j,k**). Moreover, exogenous addition of mitochondria to NHE1-overexpressing cells further exacerbates intravasation to both the trunk and the brain (**Fig. 5h,i** and **Extended Data Fig.7o**) and migration speed in the ISVs (**Fig. 5j,k**). Together, these findings indicate that activating endogenous mitochondrial function or exogenous mitochondrial transplantation can re-engage the Arp3/cytoskeleton-based migration machinery in NHE1-overexpressing cells while preserving OEM-based motility, resulting in a hyper-motile state.

## Discussion

In summary, our study uncovers a stiffness-dependent switch in cancer cell migration mechanisms and metabolic requirements under confinement from actin- and mitochondria-based motility in soft substrates to OEM- and glycolysis-dependent migration in stiff microenvironments (**Extended Data Fig. 8**). Importantly, pathophysiologically relevant mechanochemical cues, such as hypoxia and elevated extracellular fluid viscosity, are sufficient to mediate the shift from actin-based to NHE1-mediated OEM-based migration inside soft, confining spaces. This is further substantiated by the ability of cells with hyperactive NHE1 to migrate in soft microchannels and 3D collagen gels even after disruption of the actin cytoskeleton. Along these lines, the migratory defect of ILK-knockdown cells observed *in vitro* as well as in the chick embryo model, is at least partially rescued by NHE1 overexpression. In sum, our data illustrate that substrate compliance does not preclude per se OEM engagement. In this context, OEM represents a plastic migration strategy that can be engaged in compliant environments when NHE1 is sufficiently polarized or activated by cues encountered in the tumor microenvironment. Beyond cancer cells, water permeation and NHE1-dependent cell swelling were recently shown to facilitate efficient neutrophil migration in responses to chemotactic stimulation^48^. The critical role of water permeation was likewise demonstrated for motility of chemokine-activated T lymphocytes^49^.

Central to the stiffness-dependent switch is the role of NHE1 polarization and activity, regulated by YAP, which controls the transition between actin-, mitochondria-driven migration in soft microchannels and OEM-, glycolysis-driven migration in stiff ones. Consistent with this notion, NHE1 knockdown suppresses cell motility and shifts the cell metabolic dependence from glycolysis to OxPHOS in stiff substrates. Conversely, NHE1 overexpression reprograms cell bioenergetics from OxPHOS to glycolysis in soft substrates. The transition from actin-/mitochondrial-based migration to OEM-/glycolysis-driven motility may also confer an energetic advantage, as OEM engagement is associated with lower mitochondrial function and ATP/ADP ratios while preserving efficient migration. Importantly, these data illustrate that cell metabolic adaptation does not universally depend on substrate stiffness. Instead, this study provides a framework demonstrating that distinct migration mechanisms drive specific metabolic rewiring programs.

Cancer cells choose migratory programs based on energy demands, adopting strategies that minimize metabolic cost while sustaining invasion^9^. Prior work has established that amoeboid cells adopt migration behaviors that are less dependent on both adhesions and OxPHOS and have lower energetic demands compared to mesenchymal cells. This finding is conceptually consistent with our data showing that the OEM mode of migration, which is independent of β1-integrin adhesions and actin polymerization^50^, similarly operates with lower energetic demand and does not rely on OxPHOS. As such, placing the study by Crosas-Molist et al.^50^ in the context of our work, both studies support a common principle: the metabolic requirements of a migrating cell are closely coupled to the migration mechanism it employs. Along these lines, pancreatic cancer invasion in substrates of increasing stiffness is associated with increased energy demands provided by OxPHOS to support enhanced actin-/focal adhesion-dependent protrusion^16^. This prior finding is in line with our data showing that actin-based migration is supported by OxPHOS albeit in soft confining microchannels, and further highlights that cell metabolic adaptation does not universally depend on substrate stiffness. Instead, it is regulated by the cell intracellular rewiring.

The energetic optimization underlies migratory plasticity and helps explain why anti-migration therapies have been difficult to translate. By bridging OEM-driven migration to the forefront, our work reveals a fundamental, cytoskeleton-independent pathway that enables metastatic progression and exposes new therapeutic opportunities beyond traditional cytoskeletal targets. Interestingly, endogenous activation of mitochondrial function or exogenous mitochondrial transplantation into NHE1-overexpressing cancer cells results in the coexistence of both OEM-/glycolysis- and mitochondria-driven migration modes in soft microchannels. This hybrid program leads to a hypermotile cell state, observed both *in vitro* and *in vivo* in the zebrafish xenograft model. These findings highlight the plasticity of cancer cell migration strategies and suggest that cells can integrate mechanical and metabolic inputs to optimize invasiveness across diverse microenvironments.

## Materials and Experimental Methods

### Cell culture and pharmacological inhibitors

Human MDA-MB-231 (ATCC), SUM159 (ATCC), HOS cells (ATCC), and pancreatic ductal adenocarcinoma CAF-SC2 (a gift from Dr. Denis Wirtz, Johns Hopkins University) were cultured in high glucose DMEM 4.5 g/L (11995073, Gibco) with 10% Fetal Bovine Serum (FBS; 16140071, Gibco) and 1% penicillin/streptomycin (P/S; 10,000 U/mL; 15140122, Gibco). Cells were incubated at 37°C with 5% CO_2_ and 100% humidity. In select experiments cells were preconditioned for 6 days in IMDM (12440061, Life Technology Corporation)-containing media of either low (0.77 cP) or elevated (8 cP) viscosity, prepared as previously reported^18^. In other select experiments, cells were exposed to hypoxic conditions for 48 h in an InvivO_2_ 200 hypoxia workstation (Baker Ruskinn) equipped with an ICONIC digitally controlled gas mixer maintained at 1% O_2_, 5% CO_2_ and 94% N2 at 37°C.

The following drugs were used to inhibit the functions of key proteins along with corresponding vehicle controls: CK666 (100 µM; L0006, Sigma-Aldrich) for Arp2/3, CPD22 (2.5 µM; 407331, Sigma-Aldrich) for ILK, BAPTA-AM (5 µM; B6769, Invitrogen) for intracellular calcium, FTY720 (2 µM; L0700, Sigma-Aldrich) for TRPM7, Blebbistatin (20 or 5 µM; B0560, Sigma-Aldrich) for contractility, latrunculin A (LatA; 0.5 or 1 µM, Millipore Sigma) for actin, GSK 2193874 (15 µM; L0942, Sigma-Aldrich) for TRPV4, GsMTx-4 (2.5 µM; ab14871, abcam) for Piezo1/2, 2-APB (2 µM; 100065, Sigma-Aldrich) for TRP calcium channels, Rho Inhibitor I (0.5 µM; CT04-A, cytoskeleton) for RhoA activity, 2-Deoxy-D-Glucose (2-DG; 5 mM, 14325, Cayman Chemical) for glycolysis, Oligomycin (1 µM; 11341, Cayman Chemical Company) for OxPHOS, K975 (10 µM; HY138565, MedChem Express), Verteporfin (0.3 µM; SML0534, Sigma-Aldrich) for YAP, and kaempferol (1 µM; 60010, Millipore Sigma).

### Cloning, lentivirus production and cell transduction

Nontargeting scramble control, shNHE1, shAQP5, shMIIA, shMIIB, shMIIA&B, and shβ1-integrin were generated as previously described^18^. shRNA targeting ILK (5′-TCCCACGACATGCACTCAATA-3′) and MCT2 (5′-AATGTTTAGATTTGCTCAAGG-3′) were cloned into the pLKO.1 lentiviral vector (Addgene plasmid #8453, gift from B. Weinberg) using AgeI and EcoRI restriction sites. The pGP-CMV-GCaMP6s (40753; a gift from D. Kim and the GENIE Project), pLentiRhoA2G (40179; a gift from O. Pertz), and pLV-EF1a-Parkin-A92mKO2 (213559), FUGW-pHRed (65742; a gift from M. Tantama), the FUGW-PercevalHR (49083), the Peredox NADH/NAD+ biosensor (163060), and the EF1a-mito-dsRED2 (174541) plasmids were purchased from Addgene. The NHE1-GFP plasmid was a gift from J. Orlowski. For select experiments, NHE1 was amplified from the NHE1-GFP plasmid and inserted into the lentiviral backbone pLV-EF1a-IRES-Puro (Addgene, plasmid 85132). The SU9-GFP Plasmid was a gift from Dr. Sesaki^51^. MDA-MB-231, SUM159, and HOS cells were transduced as previously described^18^.

### Photolithography and PDMS device fabrication

Microfluidic devices were fabricated following previously established procedures. Photolithography was used to define microchannel architectures on silicon wafers, creating devices with multiple parallel channels. Each microchannel was designed with precise dimensions: 200 µm in length, 10 µm in height, and 3 µm in width. Polydimethylsiloxane (PDMS; Ellsworth Adhesives, #2065622) was cast onto the patterned wafer and degassed in a vacuum chamber for 30 min to eliminate air bubbles. The PDMS was then cured at 85°C for approximately 90 min. After polymerization, a 5 mm biopsy punch was used to generate six inlet and outlet wells for cell and media loading. The PDMS slabs were sonicated in 100% ethanol for 5 min, rinsed thoroughly with deionized water, and treated using a plasma cleaner (PDC-32G, Harrick) to activate the surfaces. In parallel, standard microscope glass slides (25×75 mm; Electron Microscopy Sciences, #72192-75) were cleaned with ethanol and deionized water and similarly plasma-activated. Following surface activation, PDMS and glass components were aligned and irreversibly bonded. The assembled devices were then coated with 20 µg/mL rat tail collagen type I (Gibco, A1048301) and incubated at 37°C for 1 h.

### Fabrication of polyacrylamide gels and devices

Wafers and soft microchannel devices composed of polyacrylamide (PA) were fabricated as previously reported. Solutions were prepared by combining 40% acrylamide (A) and 2% N,N′-methylenebisacrylamide (B) (both from Bio-Rad) with deionized water to achieve final concentrations of either 8% A/0.2% B or 12% A/0.25% B, corresponding to gels with stiffness values of 15 kPa and 35 kPa, respectively. Following a 20-min degassing step, polymerization was initiated by adding 10% ammonium persulfate and 0.4% TEMED (Bio-Rad). Glass coverslips (22×40 mm; Fisher Scientific) were chemically activated using glutaraldehyde as described previously. For flat gels, 60 µL of the polyacrylamide mixture was deposited onto the activated coverslip and gently flattened using a second, non-activated coverslip. For top imprinted gels, 1 mL of the polyacrylamide solution was poured over a microchannel-patterned wafer generated by photolithography and covered with a non-treated coverslip. Polymerization proceeded for 50 min at room temperature. Upon completion, gels were carefully detached: the imprinted gel was lifted off the wafer, while the flat gel remained bound to the activated coverslip. Gels were immersed in PBS without calcium or magnesium and incubated at 37 °C for 1-3 days to ensure full swelling. Once equilibrated, excess PBS was gently removed using compressed air, and inlet/outlet ports were punched into the imprinted gel. A solution of bis(sulfosuccinimidyl)suberate (BS3, 50 mg/mL; Thermo Fisher Scientific) was applied (15-20 µL) to the flat gel surface to facilitate bonding. The imprinted gel was aligned and placed onto the adhesive-coated flat gel, then incubated for 50 min at room temperature to allow crosslinking. Finally, the imprinted gel was carefully placed atop the flat gel and allowed to bond for 50 min at RT. To finalize device preparation, the assembled HEMICA devices were rehydrated in PBS (Ca2^+^/Mg2^+^-free) for 5 min. Next, 0.5 mg/mL Sulfo-SANPAH was flowed into the channels and photoactivated under UV light for 7 min. The process was repeated with fresh Sulfo-SANPAH to ensure complete activation. Finally, channels were coated with 20 µg/mL type I collagen (rat tail; Thermo Fisher Scientific) and incubated overnight at 37 °C.

### Cell seeding

Cells were detached from tissue culture dishes using 0.05% trypsin-EDTA (Gibco), centrifuged at 300 × *g* for 5 min, and resuspended at a concentration of 5 × 10⁶ cells/mL in DMEM (1X) supplemented with 1% penicillin-streptomycin and 10% heat-inactivated FBS.

For 2D migration assays, 10,000 cells per well were seeded into 24-well plates (Falcon, 3047), with or without 2D flat polyacrylamide gels^24^, which had been coated with 20 µg/mL rat tail type I collagen (Gibco) for 1.5 h at 37°C prior to each experiment. After seeding, cells were allowed to spread for 1 h, followed by drug treatment, which was maintained throughout the imaging period.

For PDMS microfluidic migration assays, 100,000 cells (5,000 cells/µL) were introduced into the seeding channel inlet, generating a pressure-driven flow across the cell seeding region. Once the flow reached the outlet, 6.67 µL of the cell suspension were aspirated from the outlet to arrest flow. Devices were then incubated at 37°C, 5% CO₂, and 100% humidity for approximately 30 min to facilitate cell spreading on the collagen I-coated seeding area near the entrances of the microchannels. Fresh media were added evenly to all six wells, and migration assays were initiated.

For PA-based microfluidic devices experiments, 10 µL of the cell suspension were added to the seeding channel inlet, generating pressure-driven flow. An equal volume of cell suspension was then added to the outlet to equilibrate the flow. Cells were allowed to adhere and spread at 37°C, 5% CO₂ for 10 min before submerging the entire device in 10 mL of complete media for subsequent migration analysis.

For experiments involving glucose or pyruvate depletion, cells were first allowed to spread as described above. Following spreading, cells were starved for 30 min in DMEM without glucose, glutamine, and pyruvate (Thermo Fisher Scientific, A1443001). After starvation, the medium was replaced with DMEM supplemented with 5% FBS and 2 mM L-glutamine (Fisher Scientific, 25030149). Depending on the experimental condition, the medium was further supplemented with 20 mM D-glucose (Agilent, 103577-100) and/or 1 mM sodium pyruvate (Agilent, 103578-100).

### PDX organoid isolation and seeding in microfluidic

All mouse husbandry and experimental procedures were conducted in accordance with protocols approved by the Johns Hopkins Medical Institute Animal Care and Use Committee (ACUC). Triple-negative breast cancer PDX (TM00096) were obtained from The Jackson Laboratory. Tumors were passaged subcutaneously in NOD.Cg-Prkdcscid Il2rgtm1Wjl/SzJ (NSG) mice for The Jackson Laboratory and were harvested prior to reaching the 20 mm endpoint. For organoid isolation, the excised tumors were mechanically minced using a scalpel and enzymatically digested using a shaker (180 rpm, 37 °C, 60 min) in collagenase solution containing collagenase (4 mg ml^−1^; C2139, Sigma-Aldrich), 5% FBS (F0926, Sigma-Aldrich), 1% penicillin-streptomycin (Sigma-Aldrich, P4333), and HEPES (15630-080, Gibco). Following digestion, samples were centrifuged at 400 xg for 10 min and treated with DNase I (2 U μl^−1^; D4263, Sigma-Aldrich). Epithelial tumor organoid clusters were enriched by differential centrifugation at 400 × g for 3 s, repeated four times, to separate epithelial clusters from stromal and single-cell components.

Harvested organoids were washed with 1x PBS and incubated in a shaker at 150 rpm and 37 °C for 10 min to dissociate the organoids into single cells. Cells were then directly seeded in collagen-coated (40 µg/ml) PDMS or PA-based µ-channels at a density of 10 million cells/mL.

### Live cell imaging

Cells were imaged every 10 min for 5-16 h on an inverted Nikon Eclipse Ti microscope (Nikon, Tokyo, Japan) equipped with automated stage and acquisition controls (NIS-Elements, Nikon) and a 10x/0.45 numerical Ph1 objective. Temperature and CO₂ levels were maintained at 37°C and 5%, respectively, using a stage-top incubator (Okolab or Tokai Hit). Cells were manually tracked in ImageJ (National Institutes of Health, Bethesda, MD) using the MTrackJ plugin. Data were analyzed in MATLAB (v. R2022b; Mathworks, Natick, MA) to determine the average velocity, speed, and persistence of each cell.

### 3D collagen migration assays

To prepare the collagen solution, 3 mL of rat tail collagen type I (354236, Corning) was mixed with 375 μL of 10× low-glucose DMEM (1 g/L; D2429, Sigma-Aldrich). The pH was gradually adjusted to physiological levels using NaOH. 25 μL of the neutralized collagen mixture was added to each well of a 24-well plate (Falcon) and incubated at 37°C for 1 h to allow gelation. Cells were detached using 0.05% trypsin-EDTA, centrifuged at 300x*g* for 5 min, and resuspended at 1×10⁶ cells/mL in DMEM (1×) supplemented with 1% penicillin-streptomycin and 10% heat-inactivated FBS. A total of 20,000 cells were mixed with 300 μL of the collagen solution, added to each well, and incubated at 37°C for 1 h to allow polymerization of the 3D collagen matrix. After gel formation, 1 mL of complete cell culture medium was gently added on top of each gel. Time-lapse imaging and migration analysis were performed as described above.

### NHE1 activity measurements using NH_4_Cl treatment

MDA-MB-231 cells were used to measure NHE1 activity under physiological and acidic pH_e_. 40,000 cells were seeded onto 24 well glass bottom dishes with or without 2D flat polyacrylamide gels^24^, coated with 20 μg/mL type I rat tail collagen, and cultured overnight. One day after seeding, cells were incubated with medium containing pHrodo^TM^ Red-AM (P35372, Molecular Probes) fluorescent dye for 30 min. The dye was then washed with 1 ml of DMEM cells were incubated with 1 mL of DMEM containing 10% FBS and 1% P/S for 10 min, and immediately imaged on a Nikon A1 confocal microscope with a Plan Apo 60x/1.4 NA objective every 30 s for 2 min. Cell culture media was then gently aspirated off and replaced with 1 mL of DMEM containing 15 mM NH_4_Cl; treated cells were imaged every 30 s for 4 min. Finally, DMEM media containing 15 mM NH_4_Cl was gently removed and replaced with 1 mL of DMEM with 10% FBS and 1% P/S; cells were imaged for every 30 s for 5 min. An ROI was manually drawn around each cell in NIS Elements Software (Nikon) to measure pHrodo^TM^ intensity values. Data were then imported into GraphPad Prism (v. 9.5.1; GraphPad, San Diego, CA) to determine the rate of pH recovery when DMEM containing 15 mM NH_4_Cl was replaced with regular DMEM. pH recovery curves were fit using simple linear regression.

### Immunofluorescence

Cells were seeded into PDMS-based devices and allowed to enter microchannels. After 24 h, cells were fixed with 4% paraformaldehyde (J19943K2, ThermoFisher Scientific) for 15 min, washed 3X with 1X PBS, and permeabilized with 1% Triton X-100 (Millipore Sigma) dissolved in 1X PBS for 10 min. Cells were then blocked for 1 h with 2% BSA and 0.1% Triton X-100 and incubated overnight at 4°C with the following primary antibodies diluted in blocking buffer: anti-NHE1 (1:50; sc-136239, Santa Cruz), anti-ILK (1:200; sc-20019, Santa Cruz), anti-β1-integrin (1:50; ITGB1 AIIB2, Developmental Studies Hybridoma Bank), anti-Arp3 (1:1000, FMS338, Abcam), or anti-TOMM20 antibody (1:250, ab186735, Abcam). Cells were washed 3X for 5 min with 1X PBS and incubated for 1 h at RT with the following secondary antibodies or fluorescent molecules diluted in 2% BSA, 0.1% Triton X-100 and 2% goat serum: Alexa Fluor 488 goat anti-rabbit (H+L) (1:200; A11034, Invitrogen), Alexa Fluor 568 goat anti-mouse (H+L) (1:1000; A11004, Invitrogen), Alexa Fluor Plus 647 goat anti-mouse (H+L) (1:100; A32728, Invitrogen), rhodamine phalloidin (1:250; R415, Invitrogen), Alexa Fluor 488 phalloidin (1:250; A12379, Invitrogen), and/or Hoechst 33342 (1:250; 561908, BD Biosciences). Finally, cells were washed 3X for 5 min with 1X PBS and imaged with a Nikon A1 or AXR confocal microscope using a Plan Apo 60x/1.4 NA (A1) or Plan Apo 40x/1.15 WI (AXR) objective, respectively. Images were analyzed in ImageJ.

### Western blotting

Western blots were carried out as previously described. Cells (200,000) were seeded into a 6 well plate (3046, Falcon) and allowed to reach 70-80% confluence. Cells were washed twice with 1 mL of ice cold 1X PBS, lysed with 60 μL of RIPA buffer (R0278, Sigma-Aldrich) containing protease/phosphatase inhibitor (5872S, Cell Signaling), scraped and collected into 1.5 mL Eppendorf tubes. After vortexing for 10 min, cell lysates were centrifugated for 10 min at 14,000g at 4°C. Protein concentrations were measured using the Pierce BCA Protein Assay (Thermo Scientific) to ensure loading of an equal amount of proteins per condition. Protein samples were boiled at 97°C for 10 min, incubated on ice for 1 min, and centrifuged for 1 min at 14,000g. Protein samples were loaded into a 4-12% Bis-Tris gel (NP0336BOX, Invitrogen) or 4-20% Tris-Glycine gel (XP04205BOX, Invitrogen); the apparatus was filled with MES SDS running buffer (NP0002, Life Technologies) or Tris/Glycine/SDS running buffer (1610732, Bio-Rad) diluted to 1X with dH_2_O; and the gel was run at 200 V for ∼50 min. Proteins were transferred from the gel onto a PVDF membrane (1704272, Bio-Rad) at 1.3 mA for 10 min. Membranes were trimmed, blocked with 5% non-fat milk for 1 h, and incubated with the following primary antibodies overnight at 4°C: anti-NHE1 (1:200; sc-136239, Santa Cruz), anti-GAPDH (14C10) (1:4000; 2118, Cell Signaling), anti-ILK (1:200; sc-20019, Santa Cruz), anti-a-tubulin (1:4000; 3873, Cell Signaling), anti-b-actin (1:4000; 4970, Cell Signaling), anti-MCT2 (1:500; 20355-1-AP, Proteintech). Membranes were then washed 5 times with 1X TBST for 5 min and incubated with the following secondary antibodies for 1 h: anti-mouse IgG HRP-linked antibody (1:2000; 7076, Cell Signaling) or anti-rabbit IgG HRP-linked Antibody (1:2000; 7074, Cell Signaling). Finally, the membranes were washed 5X with 1X TBST for 5 min, developed, and imaged.

### RNA extraction and quantitative PCR analysis

Cells (200,000) were seeded into 6 well plates and allowed to reach 80% confluence. Following two washes with 1 mL of 1× PBS, cells were lysed by adding 300 μL of TRIzol™ Reagent (15596026, Life Technologies) and incubating the lysate for 5 min at room temperature. Subsequently, 300 μL of 100% ethanol was added to each sample, and RNA was purified using the Direct-zol RNA Miniprep Plus kit (R2070, Zymo Research). Complementary DNA (cDNA) was synthesized from the isolated RNA using the iScript™ cDNA Synthesis Kit (1708891, Bio-Rad). For mitochondrial DNA (mtDNA) content quantification, total DNA was isolated using the QIAmpt DNA Mini Kit (51304, Qiagen) following the manufacturer’s instructions.

Quantitative PCR reactions were prepared in 96-well iCycler iQ plates (2239441, Bio-Rad) containing 10 μL of iTaq™ Universal SYBR Green Supermix (1725122, Bio-Rad), 7 μL of nuclease-free water, 2 μL of forward and reverse primers (20 μM), and 1 μL of cDNA template.

MCT2 forward sequence 5’-TAGCAGGAGGCTTATTATGCTGT-3’.

reverse sequence 5’- GGTTGAAGGCTAAACCTAAACCT-3’.
MT-ND1 forward sequence 5′-ATGGCCAACCTCCTACTCCT-3′.

reverse sequence 5′-CTACAACGTTGGGGCCTTT-3.
18S forward sequence 5’-CAGCCACCCGAGATTGAGCA-3’.

reverse sequence 5’-TAGTAGCGACGGGCGGTGTG-3’.

Three technical replicates were generated for each biological sample. Reactions were run on an iQ™5 Multicolor Real-Time PCR Detection System (Bio-Rad) under the following conditions: 95 °C for 30 s; 40 cycles of 95 °C for 10 s and 60 °C for 30 s; and a melt-curve stage comprising 51 increments from 70 °C to 95 °C with a 0.5 °C increase every 30 s.

### Optogenetic regulation of pAkt

Optogenetic tools were employed to control subcellular phosphorylation of Akt with high spatiotemporal precision using the Cry2-CIBN light-gated dimerization system. In this system, Cry2 is fused to the iSH2 domain of the p85α regulatory subunit of PI3K, which constitutively binds the p110α catalytic subunit, while CIBN is anchored to the plasma membrane via a CAAX motif and tagged with GFP. Upon blue light stimulation, Cry2–CIBN heterodimerization recruits the p110α subunit to the plasma membrane, inducing PI(3,4,5)P₃ formation and downstream Akt activation. The CIBN-GFP-CAAX construct was generously provided by Dr. Xavier Trepat (Institute for Bioengineering of Catalonia). The iSH2 domain was amplified from mCherry-CRY2-iSH2 (Addgene plasmid #66839) and subcloned into a lentiviral backbone (pLV-EF1α-Cry2-mCherry-puro) to generate Cry2-mCherry-iSH2 (opto-iSH2). Lentiviral transduction of MDA-MB-231 cells was followed by puromycin selection (0.5 µg/mL; A1113803, Thermo Fisher) 48 h post-transduction to establish stable expression of opto-iSH2. MDA-MB-231 cells and MDA-MB-231 NHE1-KD cells stably expressing both Cry2-mCherry-iSH2 and CIBN-GFP-CAAX were used to evaluate the role of spatiotemporally regulated Akt activation. Cells migrating in confining microchannels were imaged in real time, and their leading edges were identified by tracking mCherry fluorescence every 30 sec over a 4-min period. Optogenetic stimulation was applied to a rectangular region at the cell front using a 488 nm laser at 0.5% power for 1 sec. This stimulation was repeated every 30 sec for 16 min to maintain iSH2 enrichment at the leading edge, as confirmed by sequential mCherry imaging. Cell migration velocity was quantified using a custom MATLAB script.

### Mitochondrial Membrane Potential

Mitochondrial membrane potential was evaluated using Tetramethylrhodamine methyl ester perchlorate (TMRM; T668, Thermo Fisher Scientific) following the manufacturer’s protocol. Briefly, cells were incubated for 30 min at 37°C and 5% CO_2_ with 50 nM TMRM and subsequently seeded inside the microfluidic devices to minimize dye adsorption to the device substrate. Cells were allowed to enter the microchannels and then imaged using an AXR or NSPARC confocal microscope equipped with a Plan Apo 40×/1.15 WI objective (Nikon). TMRM was excited using a 561 nm laser line. Fluorescence intensity was quantified using ImageJ and used as a proxy for mitochondrial membrane potential.

### Total Mitochondrial content

Total mitochondrial content was assessed using MitoTracker™ Deep Red FM (M22426, Thermo Fisher Scientific). After cells had entered the microchannels or spread on 2D gels, MitoTracker was added to the culture medium at a final concentration of 1 µM. Cells were incubated for 30 min at 37°C and 5% CO₂, then washed with DMEM supplemented with 10% FBS and 1% penicillin-streptomycin before imaging. In select experiments, cells were instead washed three times with PBS, fixed with 4% PFA, and stained with Hoechst in blocking buffer. Imaging was performed using an AXR confocal microscope equipped with a Plan Apo 40×/1.15 WI objective (Nikon). MitoTracker was excited using a 640 nm laser line. Fluorescence intensity was quantified in ImageJ and used as a proxy for total mitochondrial content.

### Mitochondrial morphology analysis

Mitochondrial morphology was analyzed following previously established protocols^34^. Images were processed using the Mitochondrial Analyzer plugin in imageJ (https://github.com/AhsenChaudhry/Mitochondria-Analyzer). Background subtraction was performed using rolling radius of 1 µm, followed by application of a sigma filter plus to further reduce background noise and smooth object signal (radius = 2). Local contrast was enhanced with ‘enhance local contrast’ function (max slope = 2), and gamma was adjusted to 0.8. Mitochondria were then segmented using the weighted mean thresholding method with a block size of 1.25 µm, a C value of 5. Outliers were removed using a radius of 3 pixels.

### Exogenous addition of Mitochondria

Mitochondria were isolated from donor CAFs, as previously described^52,53^. Briefly, two million CAFs were harvested, resuspended in ice-cold mitochondrial isolation buffer (220 mM mannitol, 200 mM sucrose, 5 mM potassium monophosphate, 5 mM magnesium chloride, 1 mM EGTA, 2 mM HEPES, and 0.1% BSA in ultrapure deionized water, pH adjusted to 7.4), and homogenized on ice by passing the suspension 15-20 times through a 1 mL syringe fitted with a 27-gauge needle. The homogenate was centrifuged at 1,100 ×g for 3 min at 4 °C to remove intact cells and debris. The resulting supernatant was centrifuged at 12,000 × g for 15 min to pellet the mitochondria, which were then resuspended in 50 μL PBS and kept on ice until use. Mitochondrial protein concentration was quantified using Pierce BCA Protein Assay (Thermo Scientific). Recipient MDA-MB-231 cells were harvested, resuspended in complete media, and maintained on ice. For mitochondrial transfer, 0 or 4 μg of isolated mitochondria were added to 100,000 recipient cells, followed by centrifugation at 1500 × g for 5 min at 4 °C. For select experiments, mitochondria were isolated from mito-dsRED2-labeled CAFs, and their incorporation into unlabeled MDA-MB-231 cells was confirmed using an AXR confocal microscope equipped with a Plan Apo 20×/1.15 objective (Nikon).

### Seahorse assay

For Mitostress test, oxygen consumption rate (OCR) was measured using Seahorse extracellular flux XFPro analyzer (Agilent technologies). Following exogenous mitochondrial transplantation, cells were plated at a density of 10,000 cells/well in XFPro seahorse microplates. After 16 h, the medium was switched to Seahorse Base Medium (XF DMEM medium, #103575, Agilent technologies) supplemented with 1mM pyruvate, 10mM glucose, and 2 mM glutamine. OCR was measured following addition of 1.5 μM oligomycin, 1.5 μM FCCP and a mixture of 0.5 μM each of antimycin A and rotenone. OCR data were used to calculate the spare respiratory capacity according to the manufacturer’s instructions.

### Mitochondrial Deletion via Parkin-Mediated Mitophagy

To induce mitochondrial clearance, cells overexpressing Parkin were seeded into the desired substrate or microfluidic device and allowed to spread. Following cell spreading, 2.5 µM FCCP was added to the culture medium to depolarize mitochondrial membranes and trigger Parkin-mediated mitophagy. Cells were incubated with FCCP for 6 h at 37°C and 5% CO₂ to allow for complete mitochondrial clearance. Cells velocity was then monitored as described above. Cells were also imaged to confirm mitochondrial deletion, via mitochondrial markers (MitoTracker) as described above.

### PercevalHR ATP/ADP ratio Measurements

PercevalHR imaging was performed on a Nikon AXR confocal microscope equipped with a Plan Apo 40×/1.15 WI or 20x/1.15 objective, following established protocols^9,54^. 100,000 MDA-MB-231 cells co-expressing PercevalHR (FUGW-PercevalHR; Addgene plasmid #49083) and pHRed (FUGW-pHRed; Addgene plasmid #65742) were seeded inside the microfluidic devices in medium containing DMEM supplemented with 1% penicillin-streptomycin and 10% heat-inactivated FBS and allowed to spread and enter the microchannels. Cells were then starved for 30 min in 10 ml of phenol red-free DMEM without glucose, glutamine, and pyruvate (Thermo Fisher Scientific, A1443001). After starvation, the medium was replaced with 10 ml of phenol red-free DMEM without glucose, glutamine, and pyruvate supplemented with 5% FBS and 2 mM L-glutamine. Depending on the experimental condition, the medium was further supplemented with 20 mM D-glucose and/or 1 mM sodium pyruvate. PercevalHR was excited sequentially with 488-nm and 405-nm lasers to monitor ATP- and ADP-bound states, respectively, and emission was collected using a 500-550 nm detection window. pHRed was excited using 561-nm and 405-nm lasers, and fluorescence emission was collected with a 580-625 nm filter. PercevalHR ratio images were generated and analyzed in ImageJ using previously described workflows^9,54^.

### NADH/NAD^+^ mCherry-Peredox measurements

NADH/NAD^+^ imaging was performed on a Nikon AXR confocal microscope equipped with a Plan Apo 20x/1.15 objective. Cells expressing the Perdox mCherry biosensor were excited sequentially using 405-nm and 561-nm lasers, and emission was collected at 510-525 nm and 600-620 nm, respectively. Ratiometric 405/561 images were generated and analyzed in ImageJ using previously described workflows^55^.

### Membrane Tension and RhoA Activity Measurements

For membrane tension measurement, cells were seeded inside PA-based microchannels and allowed to enter. Cells were then treated with CPD22 or vehicle control. Next, cells were stained with live-cell membrane tension probe Flipper-TR (Spirochrome, 0.5 mM) and imaged immediately thereafter. For RhoA activity measurements, cells expressing the RhoA 2G sensor were seeded inside the mcirochannels and allowed to enter. Cells were then treated with FTY or vehicle control. FLIM analysis of live cells stained with Flipper-TR or stably expressing the RhoA2G sensor was performed as described previously using a Zeiss LSM 780 microscope and a PicoQuant system consisting of a PicoHarp 300 time-correlated single-photon counting (TCSPC) module, two-hybrid PMA-04 detectors and a Sepia II laser control module. For Flipper-TR imaging, a 485 nm laser line was used for excitation with 600/50 band-pass detector unit. For RhoA2G imaging, a 440 nm laser line was used. Cells were maintained at 37 °C and 5% CO_2_ during imaging inside a PeCon environment chamber and stage.

### Calcium Activity Measurements

Cells expressing GCAMP6s were seeded in PDMS or PA-based microchannels and allowed to spread. After spreading, cells were imaged every 30 s for 5-16 h with a 20x objective on a Nikon AXR or A1 confocal microscope using the 488 laser. The GFP signal of the cells was manually traced in ImageJ over time by measuring the mean intensity of a circular ROI inside the cell. Calcium spikes were identified as instances with greater than 2x intensity over the baseline levels.

### Total Calcium Measurements

NHE1-overexpressing cells were seeded in PA-based microchannels and allowed to enter the microchannels. Fluo-4 AM (F14201, Thermo Fisher Scientific)^24^ was then added to the culture medium at 20% (v/v). After 1 h, cells were washed twice with culture medium and immediately imaged using a 20x objective on a Nikon A1 confocal microscope using the 488 laser. Fluo-4 fluorescence intensity was quantified using ImageJ.

### Ex ovo chick embryo cancer cell extravasation model

Fertilized White Leghorn chicken eggs were obtained from the University of Alberta Poultry Research Centre and maintained in a humidified incubator at 38°C. After four days of incubation, the embryos were removed from their shells using a Dremel tool with a cutting wheel and maintained under shell-less conditions in a covered dish in a humidified air incubator at 37°C and 60% humidity, as previously described^56,57^. Cancer cell extravasation assays were performed as in^56^. Briefly, 25-50×10^3^ cancer cells were injected intravenously into the chicken CAM artery and allowed to extravasate for 6 h. 15 min before the assay CAM vasculature was visualized via injection of Lectin-649 and cancer cell extravasation was scored using intravital confocal imaging (25x objective). Random 25x imaging fields (60 μm thick; 2-3 μm step; minimum 4 fields per animal) were acquired, and individual z-stacks were analyzed to quantify the percentage of the cancer cells that extravasated out of the CAM vasculature. At least nine animals were used for each condition (three independent experiments). All the procedures were approved by the University of Alberta Institutional Animal Care and Use Committee (IACUC).

### Zebrafish husbandry and xenografts

All zebrafish (Danio rerio) procedures were conducted in accordance with NIH guidelines for the care and use of laboratory animals, and all experimental protocols with zebrafish were approved by the Johns Hopkins Institutional Animal Care and Use Committee. Zebrafish from the “AB” strain background expressing Tg(Fli1:EGFP) were crossed. Embryos were kept in E3 medium, and at 1 day post fertilization (dpf), the embryos were treated with the melanin-blocking agent PTU and maintained at 28.5 °C. At 2 or 3 dpf, the larvae were screened for EGFP expression in the vasculature using a stereo fluorescence microscope (Olympus MVX10 and/or SZX16) and manually dechorionated using tweezers. Fish from a given clutch were randomly divided into experimental groups, and all experimental groups were injected into larvae from the same clutch across at least three independent clutches. Larvae were then anesthetized with a solution of 0.4% (1x) Tricaine in E3/PTU, mounted laterally in 1% (w/v) low melting-point agarose (Fisher Scientific) on a petri dish, and immersed in the 1x Tricaine solution for the duration of the injection session. 200 mCherry-expressing MDA-MB-231 cells (100 cells/nL), with or without exogenous mitochondria addition, suspended in Hanks Balanced Saline Solution (HBSS without Ca^2+^ and Mg^2+^; Quality Biologicals; warmed to 37°C) were injected to the perivitelline space (PVS) or to the duct of cuvier (DoC) using a pulled-glass micropipette (World Precision Instruments 1B120F-4). Immediately post-injection, larvae were screened for mCherry-positive cells at the injection site via stereo fluorescence microscopy. Positive larvae were isolated and maintained overnight at 33 °C.

### Cell imaging in zebrafish

One-day after the perivitelline space injection, zebrafish larvae were anaesthetized and mounted laterally in 1% (w/v) low melting-point agarose in 24-well glass bottom plates, and immersed in the 1x Tricaine solution for the duration of the imaging. Intravasation of mCherry-labeled cancer cells to the brain and the tail was tracked via a Nikon AXR confocal microscopy using a Plan Apo 20x objective. For each larva, 3-4 confocal z-stacks centered on the vasculature, were acquired for 15 frames at 15 µm steps to image a total depth of 210 µm. The samples were simultaneously excited with a 488 and 561 nm laser and 2D images were acquired at a resolution of 1024×1024 pixels. The Nikon maximum z-projection tool was used to generate 3D images, and the number of cells intravasated into the tail and brain was quantified using the Nikon analysis software (NIS-Elements AR Analysis 5.21.03).

3 h after the duct of cuvier injection, zebrafish larvae were anaesthetized and mounted laterally in 1% (w/v) low melting-point agarose in 24-well glass bottom plates, and immersed in the 1x Tricaine solution for the duration of the imaging (∼12 h). For each larva, two confocal z-stacks centered on the vasculature, were acquired for 41 frames at 2 µm steps every 15 min. The samples were simultaneously excited with a 488 and 561 nm laser and 2D images were acquired at a resolution of 1024×1024 or 512×512 pixels. The Nikon maximum z-projection tool was used to generate 3D images, and the cells migrating through the ISVs were manually tracked in ImageJ using the MTrackJ plugin. Data were analyzed in MATLAB to determine the average speed of each cell.

### Statistical Analysis and Reproducibility

Data are presented as mean ± standard deviation (S.D.) from three or more independent biological replicates unless otherwise indicated. Individual data points represent values from single cells or experiments, as specified in the figure legends.

Normality of data distribution was assessed using the Shapiro-Wilk test for datasets with 3-8 data points, and the D’Agostino-Pearson omnibus test for datasets with more than 8 data points. For Gaussian-distributed data, statistical comparisons were made using two-tailed unpaired Student’s t-test (for two groups) or one-way ANOVA followed by Tukey’s post hoc test (for multiple groups). For log-normal distributions, data were logarithmically transformed prior to analysis using the same parametric tests. Non-Gaussian datasets were analyzed using the Mann-Whitney test (for two groups) or Kruskal-Wallis test followed by Dunn’s multiple comparisons test (for more than two groups). All statistical analyses were performed using GraphPad Prism 10 (GraphPad Software). Statistical significance was defined as p<0.05; non-significant comparisons (p>0.05) were not reported.

## Acknowledgements

The authors wish to thank Ms. Yolanda Zheng for assistance in executing select experiments.

## Author contributions

AAmitrano and DC conceptualized and designed the study, performed and analyzed many of the *in vitro* experiments, imaged and analyzed the *in vivo* zebrafish experiments, and wrote the manuscript. BI performed many *in vitro* experiments, analyzed data, and edited the manuscript. AAfthinos and SN performed select *in vitro* experiments and analyzed data and edited the manuscript. QY, BRS, and BA performed select *in vitro* experiments, analyzed data, and helped in the imaging and analysis of the *in vivo* zebrafish experiments. KS and JDL performed and analyzed the chick embryo *in vivo* experiments, provided critical input, and edited the manuscript. GG, JG, AC, and JM performed the injections for the *in vivo* zebrafish experiments, provided critical input, and edited the manuscript. MAV and SXS provided critical input and edited the manuscript. KK conceptualized, designed, supervised the study and wrote the manuscript.

## Data, Materials and Software Availability

The main data supporting the results of this study are available within the paper and its Supplementary Material file. All source data are provided with this paper. Supplementary Information is also included for this paper.

## Supplementary Information

Supplementary Fig. I: uncropped images of western blots presented in the manuscript. The red rectangles indicate the areas displayed in the figures.

**Supplementary Video 1:**

Time-lapse confocal microscopy in phase-contrast and GFP channels showing a GCAMP6 transduced MDA-MB-231 cell migrating inside PDMS-based microchannels. Time is indicated in hh:mm. Scale bar: 5 µm.

**Supplementary Video 2:**

Time-lapse confocal microscopy in phase-contrast and GFP channels showing a GCAMP6 transduced MDA-MB-231 cell migrating inside 15 kPa PA-based microchannels. Time is indicated in hh:mm. Scale bar: 5 µm.

**Supplementary Video 3:**

Time-lapse bright-field microscopy showing a Parkin-overexpressing MDA-MB-231 cell migrating inside 15 kPa microchannels after treatment with FCCP (2.5 µM). Time is indicated in hh:mm. Scale bar: 5 µm.

**Supplementary Video 4:**

Time-lapse bright-field microscopy showing a Parkin-overexpressing MDA-MB-231 cell migrating inside PDMS-based microchannels after treatment with FCCP (2.5 µM). Time is indicated in hh:mm. Scale bar: 5 µm.

**Supplementary Video 5:**

Time-lapse confocal RFP and phase-contrast showing a wild-type MDA-MB-231 cell labeled with the mitochondrial activity dye TMRM (red) migrating inside 15 kPa microchannels. Time is indicated in hh:mm. Scale bar: 5 µm.

**Supplementary Video 6:**

Time-lapse confocal microscopy in phase-contrast showing a wild-type MDA-MB-231 cell migrating inside 15 kPa microchannels in the presence of LatA (1 µM). Time is indicated in hh:mm. Scale bar: 10 µm.

**Supplementary Video 7:**

Time-lapse confocal microscopy in phase-contrast showing a wild-type MDA-MB-231 cell migrating inside 3D collagen gels in the presence of LatA (0.5 µM). Time is indicated in hh:mm. Scale bar: 100 µm.

**Supplementary Video 8:**

Time-lapse confocal microscopy in phase-contrast showing a NHE1-overexpressing MDA-MB-231 cell migrating inside 15 kPa microchannels in the presence of LatA (1 µM). Time is indicated in hh:mm. Scale bar: 10 µm.

**Supplementary Video 9:**

Time-lapse confocal microscopy in phase-contrast showing a wild-type MDA-MB-231 cell, pretreated for 6 days in 8 cP media, migrating inside 15 kPa microchannels in the presence of LatA (1 µM) in 0.77 cP media. Time is indicated in hh:mm. Scale bar: 10 µm.

**Supplementary Video 10:**

Time-lapse confocal microscopy in phase-contrast showing a NHE1-overexpressing MDA-MB-231 cell, pretreated for 6 days in 8 cP media, migrating inside 15 kPa microchannels in the presence of LatA (1 µM) in 0.77 cP media. Time is indicated in hh:mm. Scale bar: 10 µm.

**Supplementary Video 11:**

Time-lapse confocal microscopy in phase-contrast showing a NHE1-overexpressing MDA-MB-231 cell, pretreated for 6 days in 8 cP media, migrating inside 3D collagen gels in the presence of LatA (0.5 µM) in 0.77 cP media. Time is indicated in hh:mm. Scale bar: 100 µm.

---

**Extended Data Figure 1.**
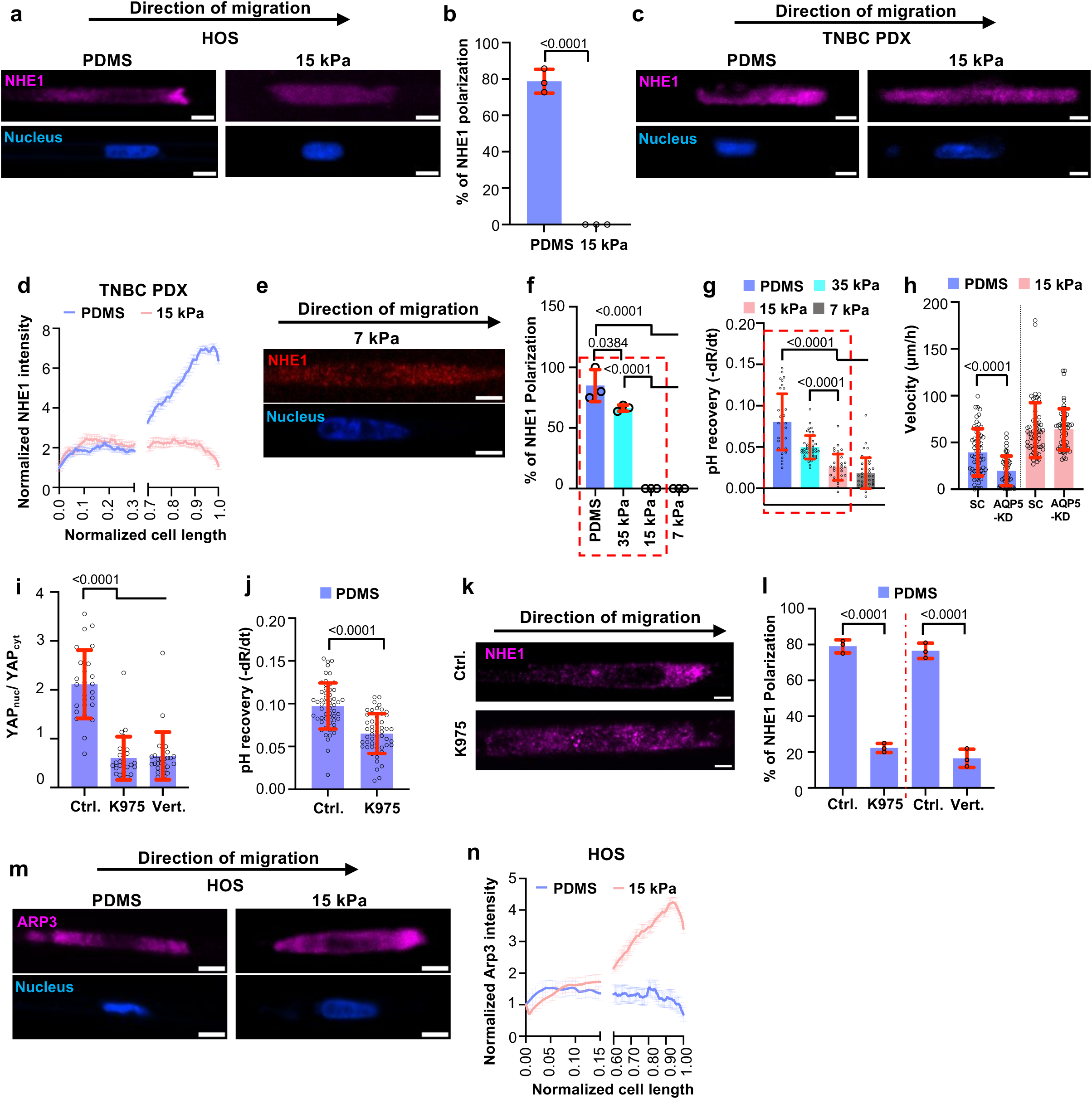
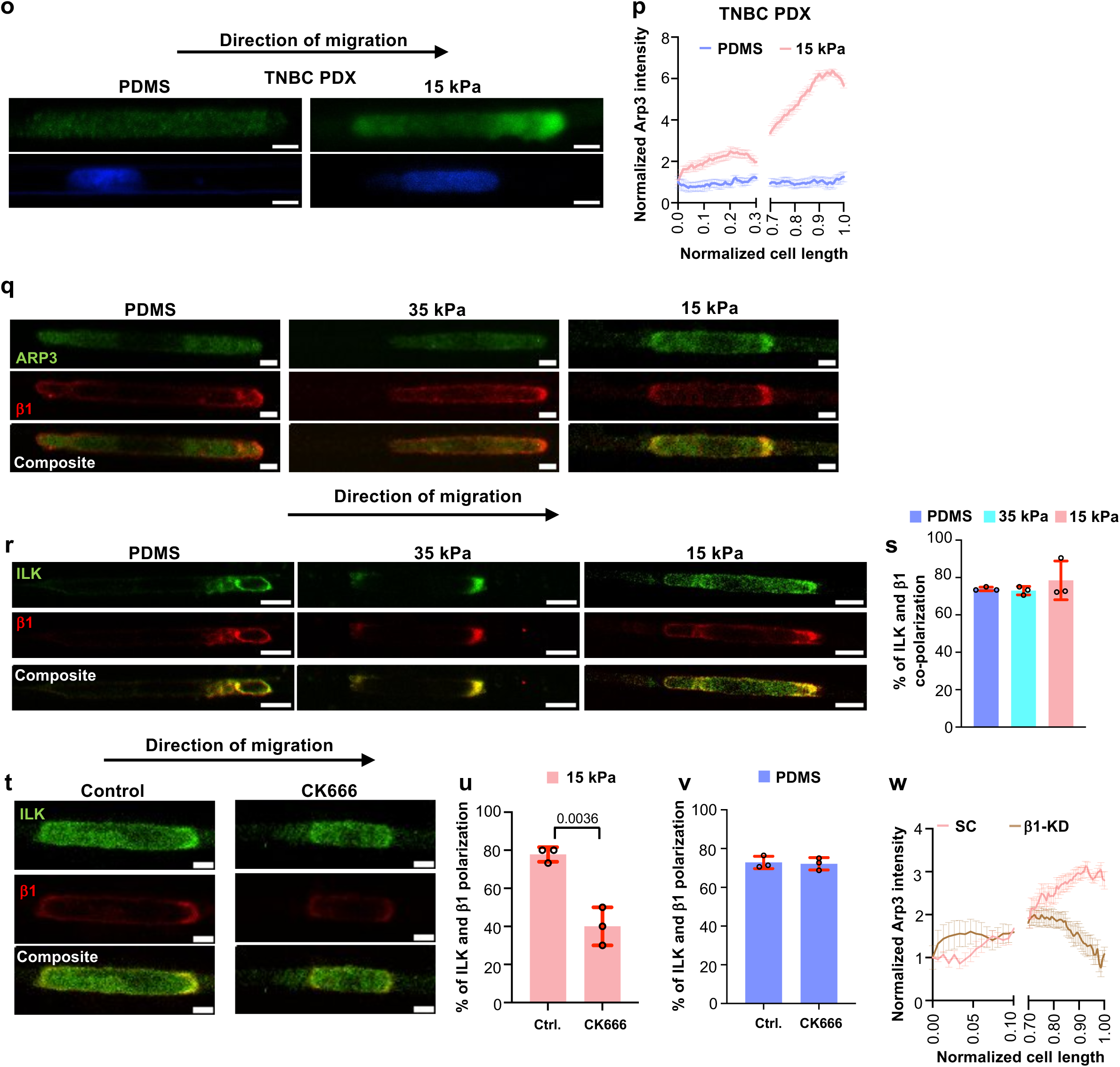
Arp3, ILK, and β1-Integrin drive migration in soft microchannels. **a**,**c,** Immunofluorescence images of HOS (a) and TNBC PDX cells (c) in PDMS- or 15 kPa PA-based collagen I-coated microchannels stained for NHE1 (magenta), and Hoechst (blue). Scale bars: 5 µm. **b**, Percentage of HOS cells in PDMS- or 15 kPa PA-based collagen I-coated microchannels displaying NHE1 polarization. Data represent mean±S.D. for n≥41 cells from 3 experiments. **d**, NHE1 intensity normalized to the cell rear and plotted along normalized cell length for TNBC PDX cells in PDMS- or 15 kPa PA-based collagen I-coated microchannels. Data are the moving average±S.D. for n≥22 cells from 2 experiments. The *x-* axis is discontinued between 0.3 and 0.7 to highlight differences at the cell edges. **e**, Immunofluorescence images of MDA-MB-231 cells in 7 kPa PA-based collagen I-coated microchannels stained for NHE1 (red), and Hoechst (blue). Scale bars: 5 µm. **f**, Percentage of MDA-MB-231 cells in PDMS- or PA-based collagen I-coated microchannels of prescribed stiffness displaying NHE1 polarization. Data represent mean±S.D. for n≥30 cells from 3 experiments. The red box indicates data displayed in Fig. 1b. **g**, Measurements of NHE1 activity of cells on 2D collagen I-coated glass or PA gels of prescribed stiffness using NH_4_Cl prepulse technique by quantifying the pH_i_ recovery of pHrodo^TM^-loaded cells. Data represent mean±S.D. for n≥36 cells from 3 experiments. The red box indicates data displayed in Fig. 1c. **h**, Migration velocity of MDA-MB-231 SC and AQP5-KD cells in PDMS-or 15 kPa PA-based collagen I-coated microchannels. Data represent mean±S.D. for n≥50 cells from 3 independent experiments. **i**, Quantification of nuclear-to-cytosolic YAP1 ratio of cells in PDMS-based microchannels in the presence of YAP1 inhibitors K975 and Verteporfin (Vert.) or vehicle control (Ctrl.; DMSO). Data represent mean±S.D. for n=27 cells from 3 independent experiments. **j,** Measurements of NHE1 activity of cells on 2D collagen I-coated glass surfaces using NH_4_Cl prepulse technique by quantifying the pH_i_ recovery of pHrodo^TM^-loaded cells. Data represent mean±S.D. for n≥53 cells from 3 experiments. **k,l,** Immunofluorescence images (k) and quantification (l) of MDA-MB-231 cells displaying NHE1 polarization in PDMS-based microchannels in the presence of YAP1 inhibitors K975 (k,l) or Verteporfin (Vert.), or vehicle control (Ctrl.; DMSO). Data represent mean±S.D. for n≥30 cells from 3 experiments. Scale bars: 5 µm. **m,o**, Immunofluorescence images of HOS (m) and TNBC PDX (o) cells in PDMS- or 15 kPa PA-based collagen I-coated microchannels stained for Arp3 (magenta (m) or green (o)) and Hoechst (blue). Scale bars: 5 µm. **n**,**p,** ARP3 intensity normalized to the cell rear and plotted along normalized cell length for HOS (n) and TNBC PDX (p) in PDMS- or 15 kPa PA-based collagen I-coated microchannels. Data are the moving average±S.D. for n≥26 cells from 3 (n) or 2 (p) experiments. The *x-* axis is discontinued between 0.15 and 0.6 (n) or 0.3 and 0.7 (p) to highlight differences at the cell edges. **q,** Immunofluorescence images of cells in PDMS- or PA-based collagen I-coated microchannels stained for Arp3 (green) and β1-integrin (red). Scale bars: 5 µm. **r,s**, Immunofluorescence images (r) and percentage of MDA-MB-231 cells (s) in PDMS- or PA-based collagen I-coated microchannels displaying ILK (green) and β1-integrin (red) front polarization. Data represent mean±S.D. for n≥30 cells from 3 independent experiments. Scale bars: 5 µm. **t,u**, Immunofluorescence images (t) and percentage of MDA-MB-231 cells (u) in 15 kPa PA-based collagen I-coated microchannels displaying ILK (green) and β1-integrin (red) front polarization in the presence of Arp2/3 inhibitor CK666 or vehicle control (Ctrl.; DMSO). Data represent mean±S.D. for n≥30 cells from 3 experiments. Scale bars: 5 µm. **v**, Percentage of MDA-MB-231 cells in PDMS-based collagen I-coated microchannels displaying ILK and β1-integrin front polarization in the presence of Arp2/3 inhibitor CK666 or vehicle control (Ctrl.; DMSO). Data represent mean±S.D. for n≥43 cells from 3 experiments. **w**, Arp3 intensity normalized to the cell rear and plotted along normalized cell length in confined MDA-MB-231 SC or β1-integrin-KD cells. Data are the moving average±S.D. for n≥24 cells from 3 experiments. The *x-* axis is discontinued between 0.10 and 0.70 to highlight differences at the cell edges. Statistical significance was assessed by unpaired t-test (b, i, l), one-way ANOVA followed by Tukey’s (f), Kruskal-Wallis (g,i), Mann-Whitney (l,u).

**Extended Data Figure 2.**
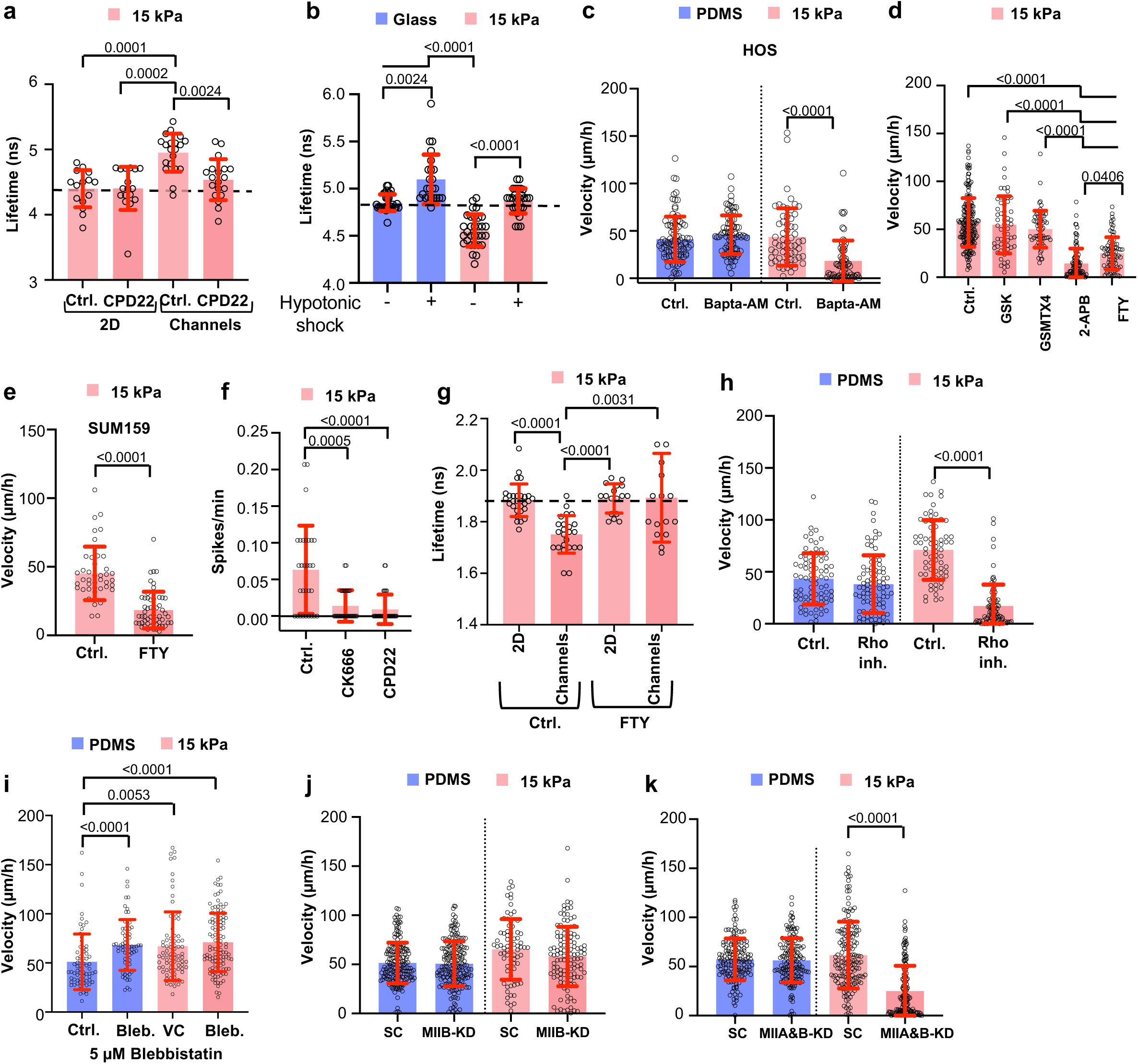
Cells use distinct migratory programs in soft versus stiff microchannels. **a**, Flipper-TR lifetime quantification of membrane tension of MDA-MB-231 cells in 15 kPa PA-based collagen I-coated surfaces in the presence of ILK inhibitor CPD22 or vehicle control (Ctrl.; DMSO). Data are the moving average±S.D. for n≥20 cells from 3 experiments. **b**, Flipper-TR lifetime quantification of membrane tension of MDA-MB-231 cells in 2D glass or 15 kPa PA-based collagen I-coated surfaces in the presence or absence of hypotonic shock. Data are the moving average±S.D. for n≥29 cells from 3 experiments. **c**, Migration velocity of HOS cells in PDMS- or 15 kPa PA-based collagen I-coated microchannels in the presence of the calcium chelator BAPTA-AM or vehicle control (Ctrl.; DMSO). Data represent mean±S.D. for n≥60 cells from 3 independent experiments. **d**, Migration velocity of MDA-MB-231 cells in 15 kPa PA-based collagen I-coated microchannels in the presence of the TRPV4 inhibitor GSK, or the Piezo1 inhibitor GSMTX4, or the TRP channels inhibitor 2-APB, or the TRPM7 inhibitor FTY720, or vehicle control (Ctrl.; DMSO). Data represent mean±S.D. for n≥61 cells from 3 independent experiments. **e**, Migration velocity of SUM-159 cells in 15 kPa PA-based collagen I-coated microchannels in the presence of the TRPM7 inhibitor FTY720 or vehicle control (Ctrl.; DMSO). Data represent mean±S.D. for n≥42 cells from 3 experiments. **f**, GCaMP6 spikes/min of MDA-MB-23cells in 15 kPa PA-based collagen I-coated microchannels in the presence of ARP2/3 inhibitor CK666, ILK inhibitor CPD22, or vehicle control (Ctrl.; DMSO). Data are mean±S.D. for n=30 cells from 3 experiments. **g**, Lifetime of RhoA activity biosensor in MDA-MB-231 RhoA2G-cells in 15 kPa PA-based collagen I-coated 2D area and microchannels in the presence of TRPM7 inhibitor FTY720 or vehicle control (Ctrl.; DMSO). Data are mean±S.D. for n≥16 cells from 3 experiments. **h**, Migration velocity of MDA-MB-231 cells in PDMS- or 15 kPa PA-based collagen I-coated µ-channels in the presence of the RhoA inhibitor or vehicle control (Ctrl.; DMSO). Data represent mean±S.D. for n≥69 cells from 3 experiments. **i**, Migration velocity of MDA-MB-231 cells in PDMS- or 15 kPa PA-based collagen I-coated microchannels in the presence of blebbistatin or vehicle control (Ctrl.; DMSO). Data represent mean±S.D. for n≥71 cells from 3 independent experiments. **j**, Migration velocity of MDA-MB-231 SC or MIIB-KD cells in PDMS- or 15 kPa PA-based collagen I-coated microchannels. Data represent mean±S.D. for n≥74 cells from 3 experiments. **k**, Migration velocity of MDA-MB-231 SC or MIIA and MIIB dual KD (MIIA&B-KD) cells in PDMS-or 15 kPa PA-based collagen I-coated microchannels. Data represent mean±S.D. for n≥158 cells from 3 experiments. Statistical significance was assessed by Kruskal-Wallis (a,b,d,f,g,i), Mann-Whitney (c,h,k), unpaired t-test (e).

**Extended Data Figure 3.**
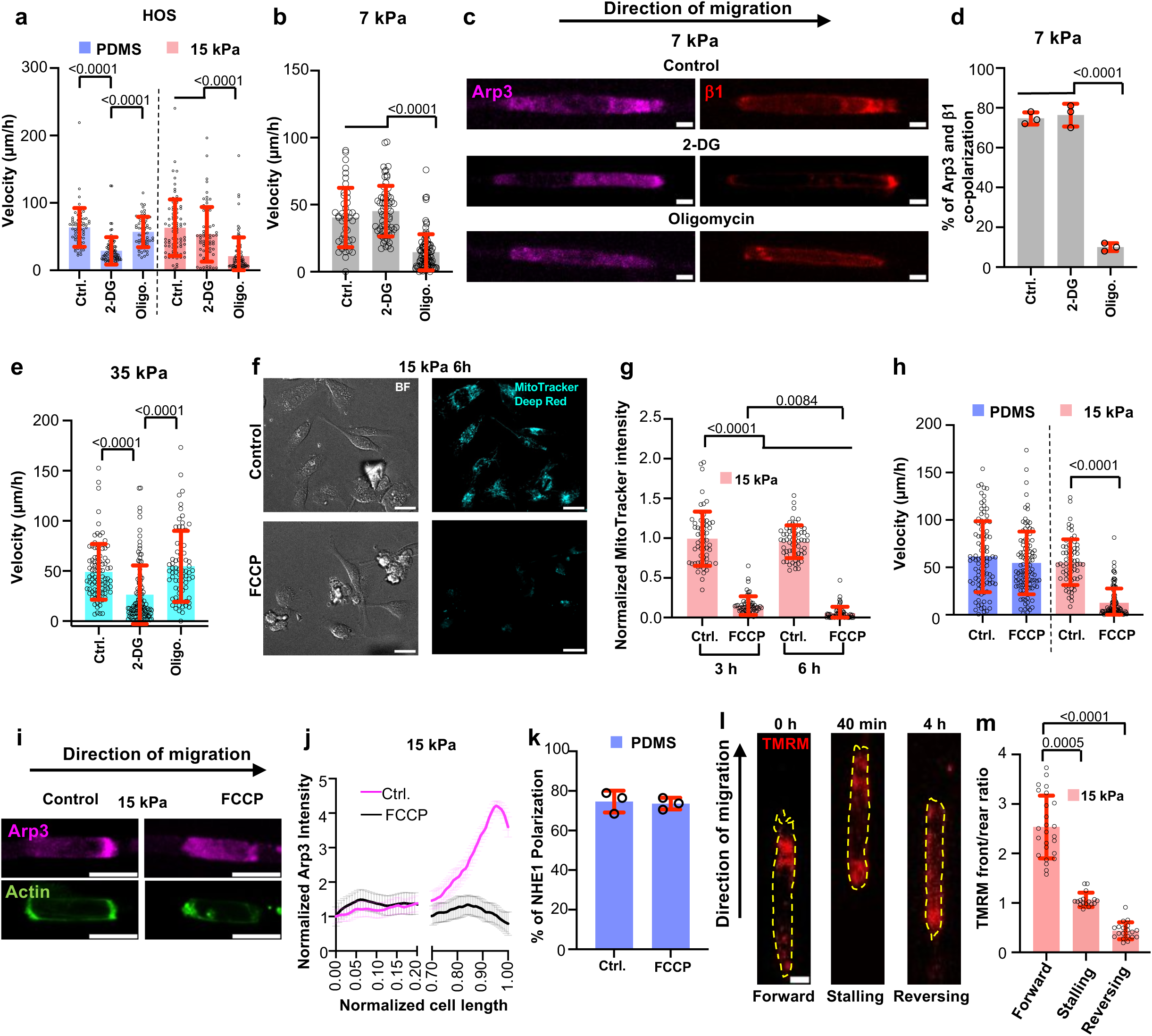
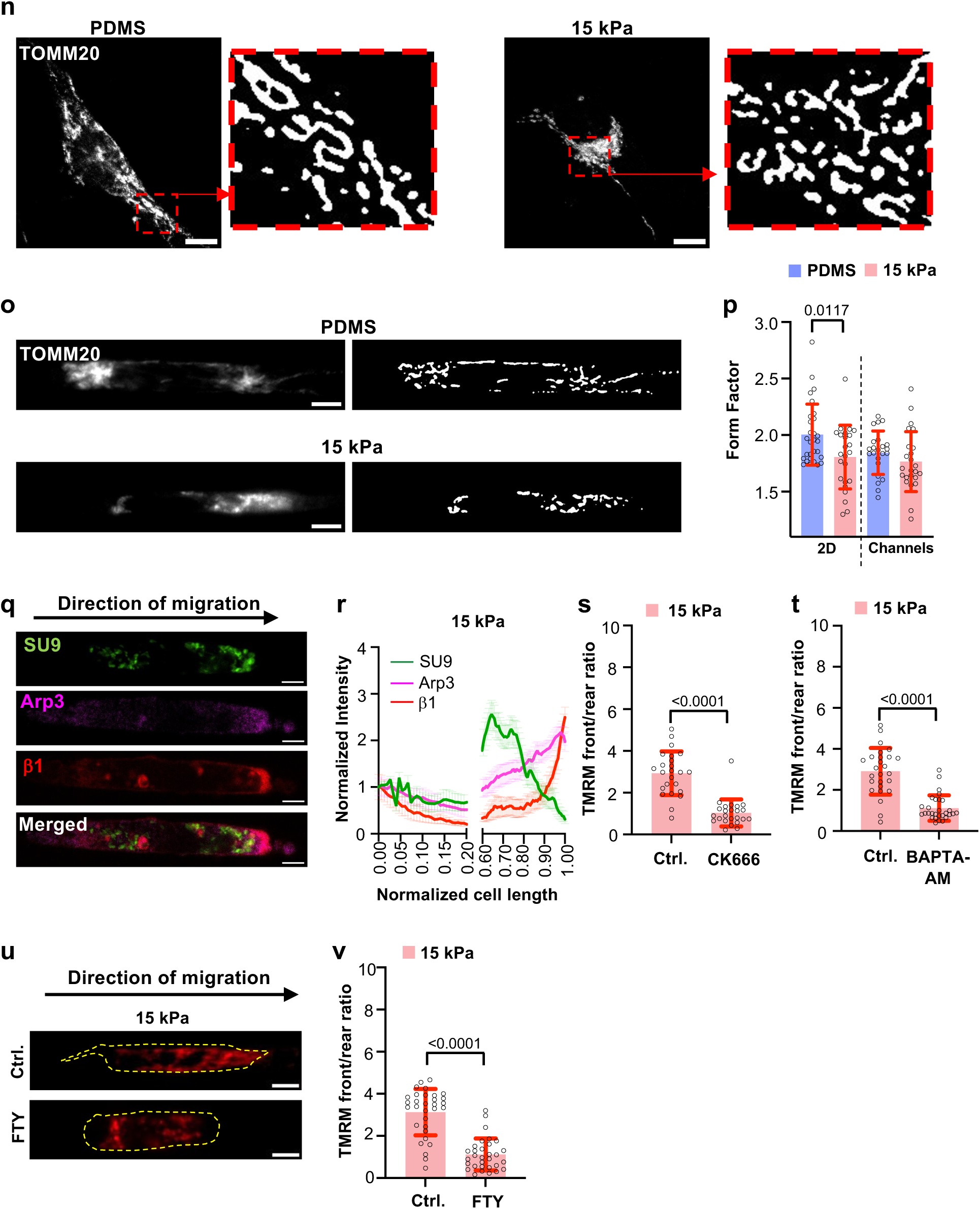
Distinct metabolic requirements for cell motility in soft versus stiff microchannels. **a**, Migration velocity of HOS cells in PDMS- or 15 kPa PA-based collagen I-coated microchannels in the presence of 2-DG or oligomycin (Oligo.) or vehicle control (Ctrl.; DMSO). Data represent mean±S.D. for n≥68 cells from 3 experiments. **b**,**e**, Migration velocity of MDA-MB-231 cells in 7 (b) or 35 (e) kPa PA-based collagen I-coated microchannels in the presence of 2-DG or oligomycin (Oligo.) or vehicle control (Ctrl.; DMSO). Data represent mean±S.D. for n≥66 cells from 3 experiments. **c**, Immunofluorescence images of cells in 7 kPa PA-based collagen I-coated microchannels stained for Arp3 (magenta) and β1-integrin (red) in the presence of 2-DG or oligomycin or vehicle control (Ctrl.; DMSO). Scale bars: 5 µm. **d**, Percentage of MDA-MB-231 cells displaying front Arp3 and β1-integrin co-polarization in 7 kPa PA-based collagen I-coated microchannels in the presence of 2-DG or Oligomycin (Oligo.) or vehicle control (Ctrl.; DMSO). Data are mean±S.D. from n=30 cells from 3 experiments. **f**,**g**, Representative immunofluorescence images (f) and normalized MitoTracker intensity (g) of MDA-MB-231 cells on 15 kPa PA-based collagen I-coated 2D surfaces in the presence of FCCP or vehicle control (Ctrl.; DMSO). Data are mean±S.D. for n=61 cells from 3 experiments. Scale bars: 20 µm. **h**, Migration velocity of Parkin overexpressing cells in PDMS- or 15 kPa PA-based collagen I-coated microchannels in the presence of FCCP or vehicle control (Ctrl.; DMSO). Data represent mean±S.D. for n≥65 cells from 3 experiments. **i**,**j**, Representative immunofluorescence images (i) and Arp3 intensity normalized to the cell rear and plotted along the normalized cell length (j) in Parkin overexpressing cells in 15 kPa PA-based collagen I-coated microchannels in the presence of FCCP or vehicle control (Ctrl.; DMSO) stained for Arp3 (magenta) and phalloidin (green). Scale bars: 10 µm. **k**, Percentage of Parkin overexpressing MDA-MB-231 cells displaying front NHE1 polarization in PDMS collagen I-coated microchannels in the presence of FCCP or vehicle control (Ctrl.; DMSO). Data are mean±S.D. for n≥53 cells from 3 experiments. **l**,**m**, Representative immunofluorescence images (l) and corresponding front/rear ratios (m) of TMRM (red) intensity in cells migrating in 15 kPa PA-based collagen I-coated microchannels, shown during forward movement, stalling, or reversal. Scale bars: 5 µm. Data are mean±S.D. for n=61 cells from 3 experiments. **n,o,** Representative images of TOMM20 staining and corresponding thresholded images of cells on 2D surfaces (n) or within microchannels of PDMS- or 15 kPa PA-based microfluidic devices. Scale bars: 5 µm. **p,** Quantification of mitochondrial form factor of cells on 2D surfaces or within microchannels of PDMS- or 15 kPa PA-based microfluidic devices. Data are mean±S.D. for n≥23 cells from 3 experiments. **q**,**r**, Immunofluorescence images (q) and intensity normalized to the cell rear and plotted along the normalized cell length (r) for SU9-GFP cells in 15 kPa PA-based collagen I-coated microchannels stained for Arp3 (magenta) and β1-integrin (red). Scale bars: 5 µm. Data are mean±S.D. for n≥81 cells from 3 experiments. The *x-* axis is discontinued between 0.20 and 0.60 to highlight differences at the cell edges. **s,t**, Cell front/rear TMRM ratio in 15 kPa PA-based collagen I-coated microchannels in the presence of Arp2/3 inhibitor CK666 (s) or calcium chelator BAPTA-AM (t) or vehicle control (Ctrl.; DMSO). Data represent mean±S.D. for n≥24 cells from 3 experiments. **u**,**v**, Representative immunofluorescence images (u) and front/rear ratio (v) of TMRM (red) intensity in cells in 15 kPa PA-based collagen I-coated microchannels in the presence of TRPM7 inhibitor FTY720 or vehicle control (Ctrl.; DMSO). Scale bars: 5 µm Data represent mean±S.D. for n=31 cells from 3 experiments. Statistical significance was assessed by Kruskal-Wallis (a,b,e,g,m), one-way ANOVA followed by Tukey’s (d), unpaired t-test (h,p) after lognormal transformation (s,v) or Mann-Whitney (t).

**Extended Data Figure 4.**
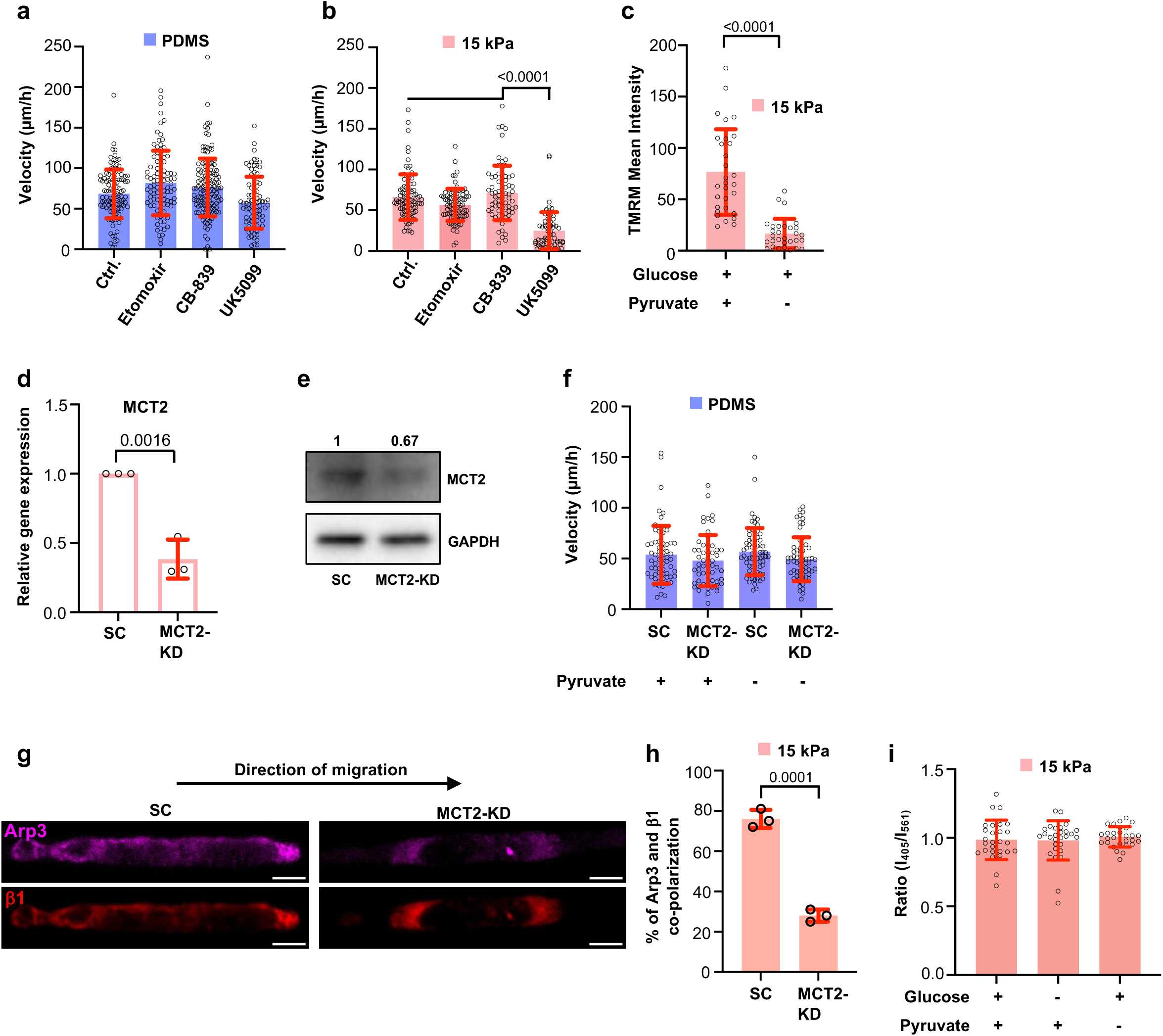
MCT2-driven pyruvate uptake fuels mitochondria-based migration in soft microchannels. **a**,**b**, Cell migration velocity in PDMS- (a) or 15 kPa PA-based (b) collagen I-coated microchannels in the presence of fatty acid oxidation inhibitor Etomoxir, glutaminase inhibitor CB-839, mitochondrial pyruvate carrier inhibitor UK-5099 or vehicle control (Ctrl.; DMSO). Data represent mean±S.D. for n≥93 cells from 3 experiments. Statistical significance was assessed by Kruskal-Wallis. **c**, Mean TMRM intensity for cells in 15 kPa PA-based collagen I-coated microchannels in the presence or absence of pyruvate. Data represent mean±S.D. for n=30 cells. Statistical significance was assessed by Mann-Whitney. **d**, Quantification of MCT2 mRNA expression of SC and MCT2-KD cells. Data represent mean±S.D. from 3 experiments. Statistical significance was assessed by unpaired t-test. **e**, Representative WB image of MCT2 protein expression in SC and MCT2-KD cells. **f**, Migration velocity of SC or MCT2-KD cells in PDMS-based collagen I-coated microchannels in medium with or without pyruvate. Data represent mean±S.D. for n≥61 cells from 3 independent experiments. **g**,**h,** Immunofluorescence images (g) and quantification (h) of SC and MCT2-KD cells displaying Arp3 (magenta) and β1-integrin (red) front polarization. Data represent mean±S.D. for n=30 from 3 experiments. Scale bars: 5 µm. Statistical significance was assessed by unpaired t-test. **i**, Ratiometric pH values (I_405_/I_561_) in cells co-expressing Perceval HR and pHRed inside 15 kPa PA-based microchannels in medium with or without glucose or pyruvate. Data represent mean±S.D for n≥27 cells from 3 experiments. Cell model: MDA-MB-231.

**Extended Data Figure 5.**
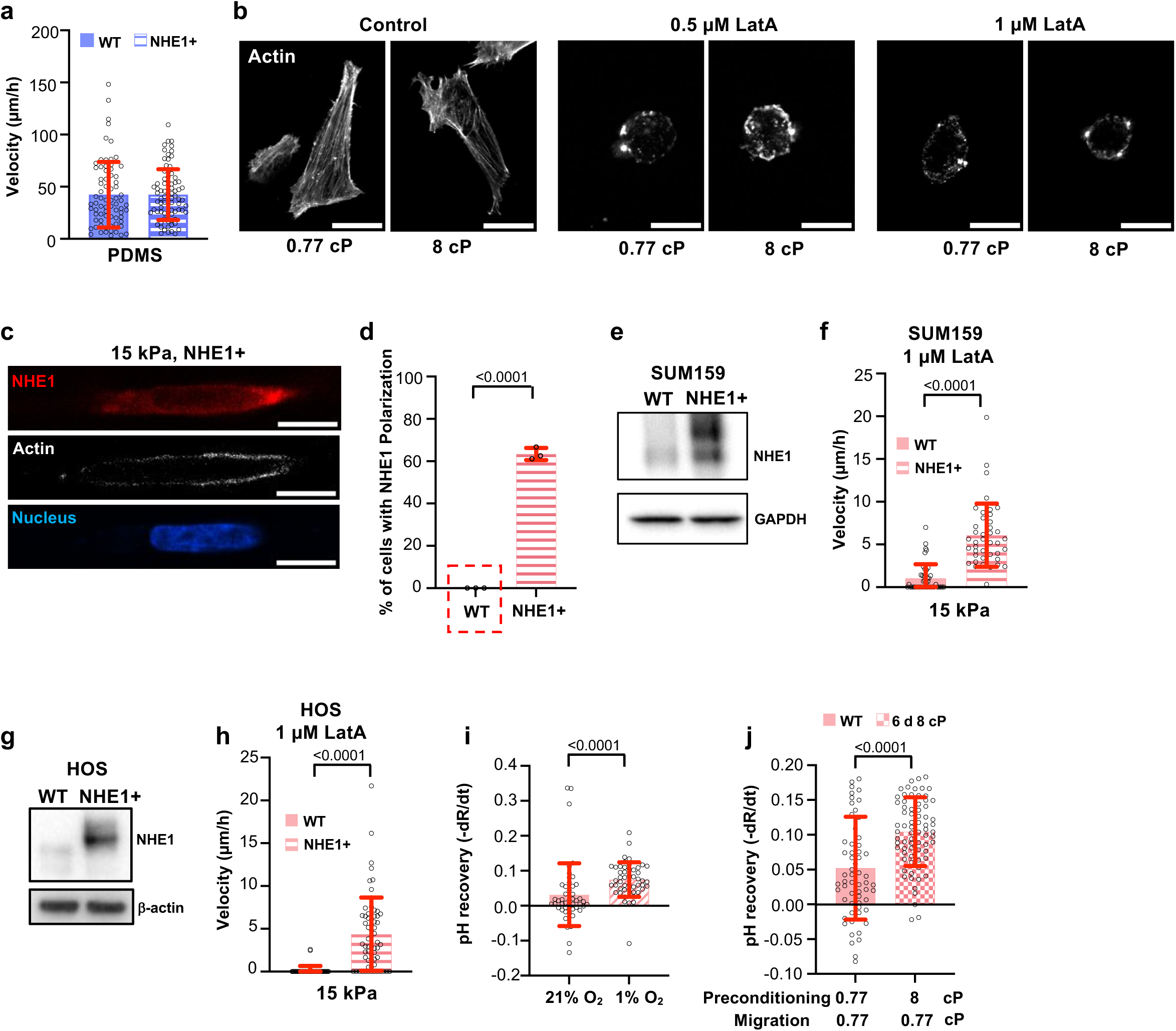
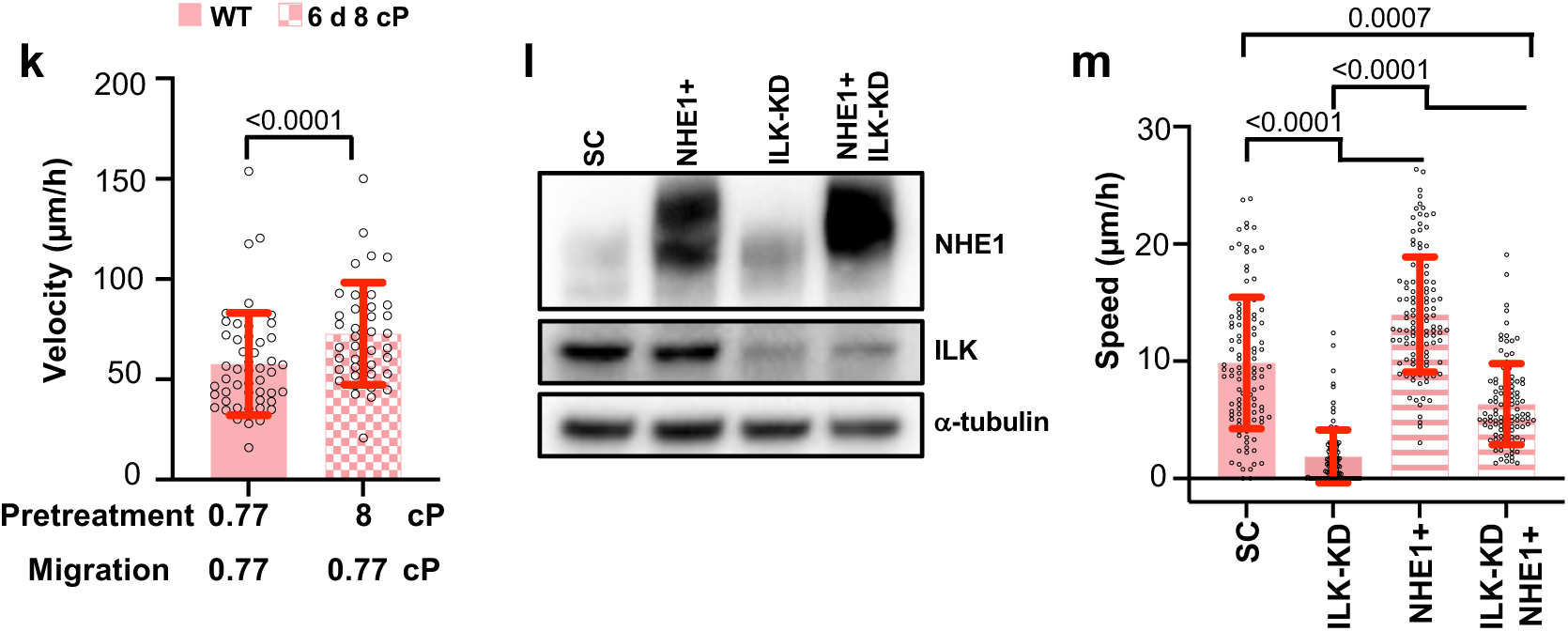
NHE1 overexpression and activation promotes OEM-based migration in soft microchannels. **a**, Migration velocity of MDA-MB-231 wild-type and NHE1-overexpressing cells in PDMS-based collagen I-coated microchannels. Data are mean±S.D. for n≥73 cells from 3 experiments. **b**, Representative images of MDA-MB-231 cells on 2D collagen I-coated 15 kPa PA gels and treated with LatA (0.5 or 1 µM) or vehicle control (DMSO) for 2 h before fixing and staining them with phalloidin (white). **c**,**d**, Representative immunofluorescence image (c) and percentage of cells displaying NHE1 polarization (d) in NHE1-overexpressing cells in 15 kPa PA-based collagen I-coated microchannels. Data are mean±S.D. from 3 experiments. Scale bars: 10 µm. The red box indicates data displayed in Fig. 1b. **e**,**g**, Representative WB image of NHE1 protein expression in SUM-159 (e) and HOS (g) wild-type and NHE1-overexpressing cells. **f**,**h**, Migration velocity of SUM-159 (f) and HOS (h) wild-type and NHE1-overexpressing cells in 15 kPa PA-based collagen I-coated microchannels in the presence of LatA. Data are mean±S.D. for n≥42 cells from 3 experiments. **i**, Measurements of NHE1 activity of MDA-MB-231 cells preconditioned to normoxia or hypoxia for 48 h on 2D collagen I-coated 15 kPa PA gels using NH_4_Cl prepulse technique by quantifying the pH_i_ recovery of pHrodo^TM^-loaded cells. Data represent mean±S.D. for n=48 cells from 3 experiments. **j**, Measurements of NHE1 activity of MDA-MB-231 cells preconditioned for 6 days at 0.77 or 8 cP and then seeded on 2D collagen I-coated 15 kPa PA gels using NH_4_Cl prepulse technique by quantifying the pH_i_ recovery of pHrodo^TM^-loaded cells. Data represent mean±S.D. for n≥61 cells from 3 experiments. **k**, Migration velocity of MDA-MB-231 cells preconditioned for 6 days at 0.77 or 8 cP and then seeded in 15 kPa PA-based collagen I-coated microchannels. Data represent mean±S.D. for n≥50 cells from 3 experiments. **l**, Representative WB image of NHE1 and ILK protein expression in MDA-MB-231 SC, NHE1-overexpressing, ILK-KD, and NHE1-overexpressing/ILK-KD cells. **m**, Migration speed of MDA-MB-231 SC, NHE1-overexpressing, ILK-KD, and ILK-KD NHE1-overexpressing in 3D collagen. Data are the mean ± S.D. for n≥102 cells from 3 experiments. Statistical significance was assessed by unpaired t-test (d,j) after lognormal transformation (k), Mann-Whitney (f,h,i), or Kruskal-Wallis (m).

**Extended Data Figure 6.**
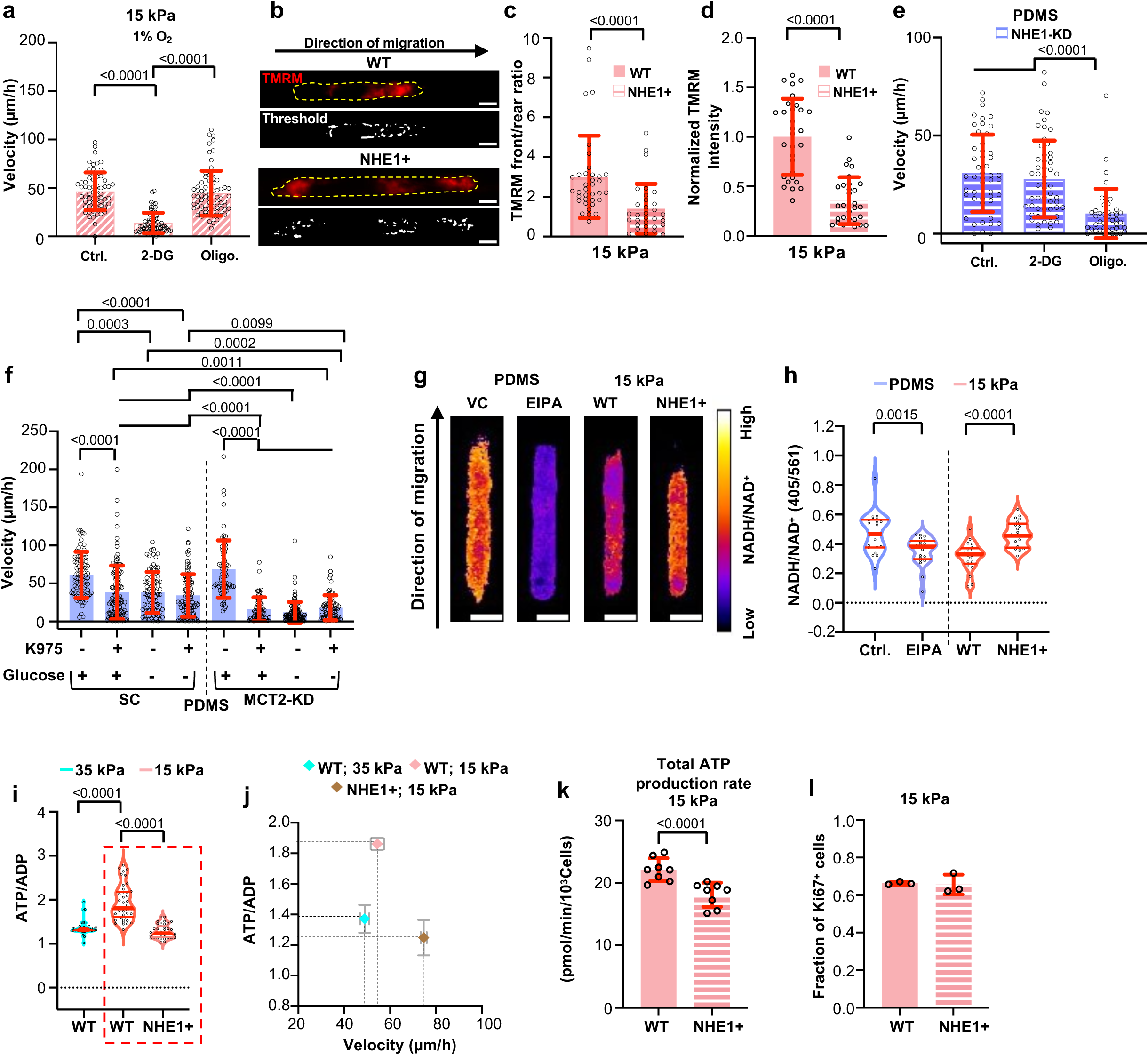
Cell intracellular rewiring overrides stiffness-dependent metabolic adaptation. **a,** Cell migration velocity after exposure to hypoxia for 48 h and seeded in 15 kPa PA-based collagen I-coated microchannels in the presence of 2-DG or oligomycin (oligo.) or vehicle control (Ctrl.; DMSO). Data represent mean±S.D. for n≥66 cells from 3 experiments. **b-d**, Representative immunofluorescence and thresholded images (b) front/rear ratio (c) and normalized intensity (d) of TMRM (red) in wild-type or NHE1-overexpressing cells in 15 kPa PA-based collagen I-coated microchannels. Cell perimeter is denoted by the yellow lines. Data represent mean±S.D. for n≥30 cells from 3 experiments. Scale bars: 5 µm. **e**, Migration velocity of NHE1-KD cells in PDMS-based collagen I-coated microchannels in the presence of 2-DG or oligomycin (Oligo.) or vehicle control (Ctrl.; DMSO). Data represent mean±S.D. for n≥44 cells from 3 experiments. **f**, Migration velocity of SC or MCT2-KD cells in PDMS-based collagen I-coated microchannels in media with or without glucose in the presence or absence of YAP1 inhibitor K975. Data represent mean±S.D. for n≥65 cells from 3 experiments. **g**,**h**, Representative immunofluorescence images (g) and quantification (h) of 405/561 ratio of wild-type or NHE1-overexpressing cells carrying the NADH/NAD^+^ Peredox biosensor in PDMS- or 15 kPa PA-based microchannels in the presence of EIPA or vehicle control (Ctrl.; DMSO). Data represent mean±S.D. for n=20 cells from 3 experiments. Red lines indicate the median (thick) and quartile (thin). Scale bars: 5 µm. **i**, Quantification of PercevalHR (ATP/ADP) ratio of wild-type or NHE1-overexpressing cells co-expressing Perceval HR and pHred in 35 or 15 kPa PA-based microchannels. Data represent mean±S.D. for n≥39 cells from 3 independent experiments. Red lines indicate the median (thick) and quartile (thin). Scale bars: 5 µm. The red box indicates data displayed in Fig. 5c. **j**, ATP/ADP ratio as a function of migration velocity in WT and NHE1-overexpressing cells migrating in 35 or 15 kPa microchannels. Data are mean±S.D. for n≥22 cells from 3 experiments. **k**, Quantification of ATP production rate of WT and NHE1-overexpressing cells on 15 kPa 2D gels via Seahorse Analyzer. Data represent mean±S.D. from 3 experiments. **l**, Fraction of Ki-67 positive WT and NHE1-overexpressing cells on 15 kPa 2D gels. Data represent mean±S.D. from 3 experiments. Statistical significance was assessed by Kruskal-Wallis (a,e,f,i), Mann-Whitney (d), unpaired t-test (h,k) after log transformation (c). Cell model: MDA-MB-231.

**Extended Data Figure 7.**
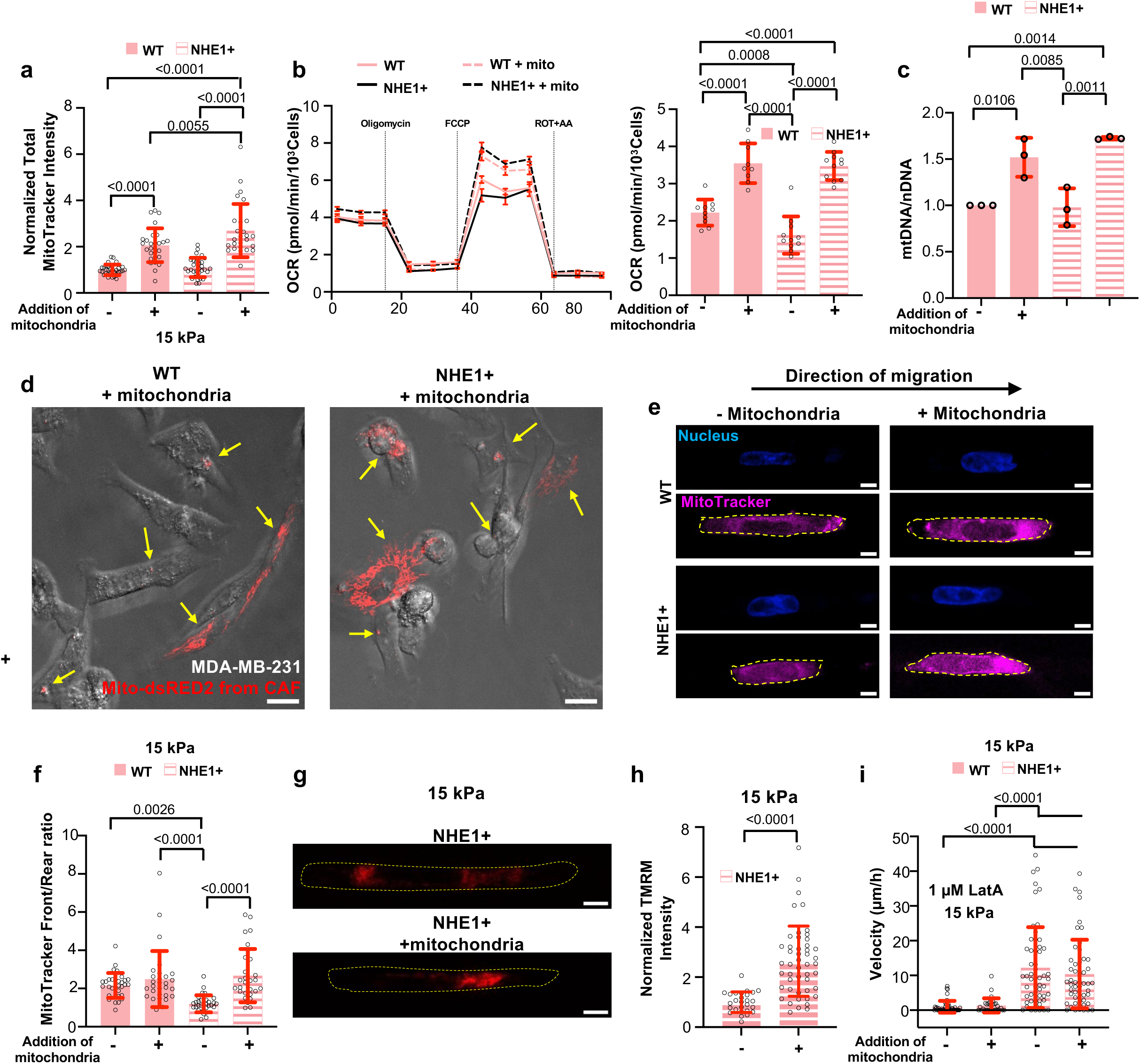
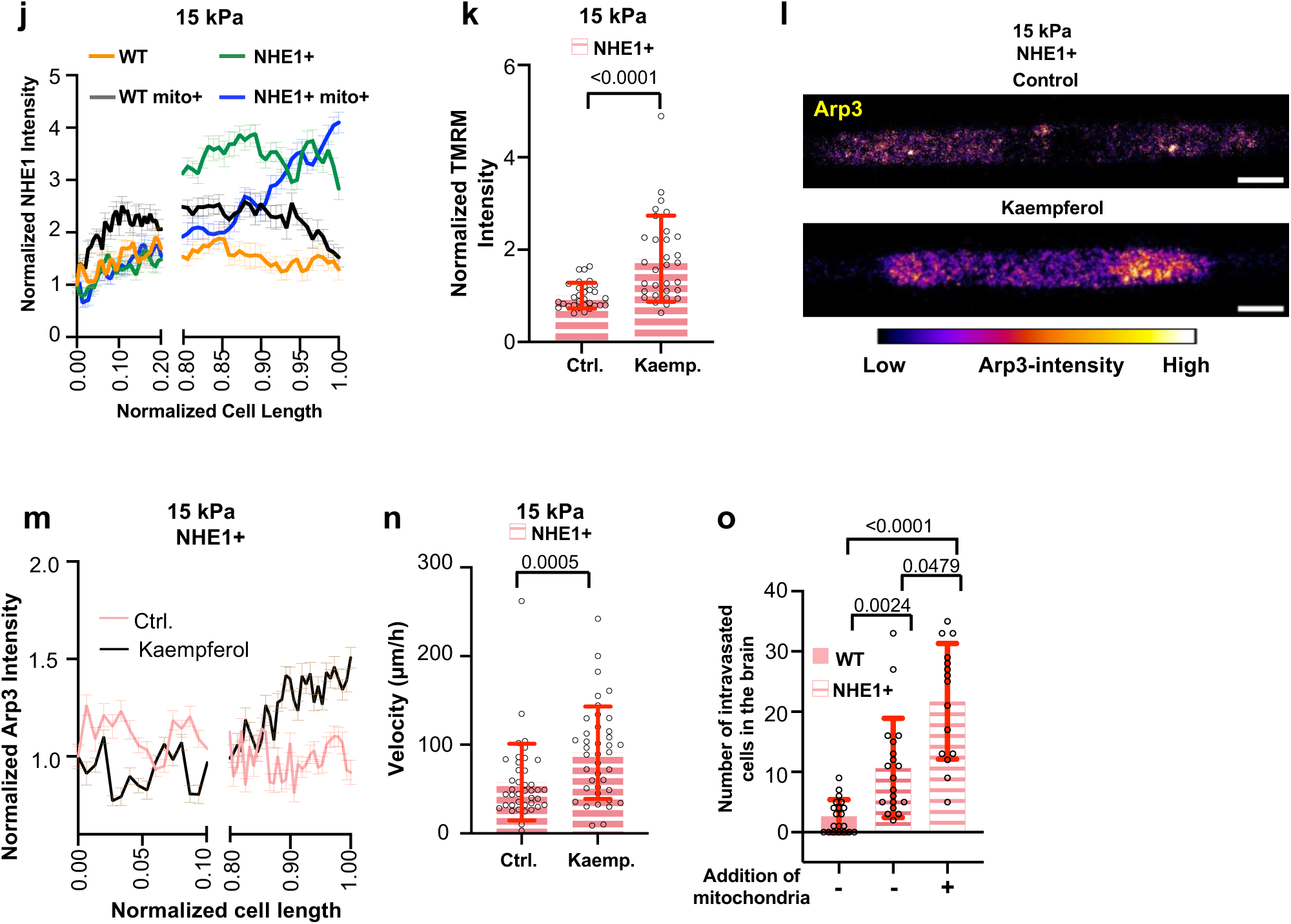
Mitochondrial activation promotes co-engagement of migration modes resulting in a hypermotile state. **a**, Normalized overall MitoTracker intensity in wild-type or NHE1-overexpressing cells in 15 kPa PA-based collagen I-coated microchannels with or without exogenous mitochondrial addition. Data represent mean±S.D. for n≥29 cells from 3 experiments. **b**, Mitochondrial spare respiratory capacity (right) measured from oxygen consumption rate (OCR) (left) of wild-type and NHE1-overexpressing cells with or without exogenous mitochondrial addition. Data represent mean±S.D. from 2 experiments. **c**, Quantitative real-time PCR quantification of mtDNA relative to nDNA (mtDNA/nDNA) as measured by primers specific to mitochondrial gene MT-ND1 and nuclear gene 18s for WT or NHE1-overexpressing cells with or without mitochondrial addition. Data represent mean±S.D. from 3 experiments. **d**, Representative immunofluorescence images showing successful incorporation of mitochondria isolated from Mito-dsRED2-labeled CAFs (red) into unlabeled recipient WT or NHE1-overexpressing cells (brightfield). Yellow arrows point to mitochondria from CAFs. Scale bars: 20 µm. **e**,**f**, Representative immunofluorescence images (e) and front/rear ratio (f) of MitoTracker (magenta) intensity in wild-type or NHE1-overexpressing cells in 15 kPa PA-based collagen I-coated microchannels. Data represent mean±S.D. for n≥29 cells from 3 experiments. Scale bars: 5 µm. **g,h,** Representative immunofluorescence images (g) and normalized intensity (h) of TMRM in NHE1-overexpressing cells in 15 kPa PA-based collagen I-coated microchannels. Data represent mean±S.D. for n≥27 cells from 3 experiments. Scale bars: 5 µm **i**, Migration velocity of wild-type and NHE1-overexpressing cells in 15 kPa PA-based collagen I-coated microchannels with or without exogenous mitochondrial addition and treated with LatA. Data represent mean±S.D. for n≥33 cells from 3 experiments. **j**, NHE1 intensity normalized to the cell rear and plotted along the normalized cell length in wild-type and NHE1-overexpressing cells in 15 kPa PA-based collagen I-coated microchannels with or without exogenous mitochondrial addition. Data are the moving average±S.D. for n≥17 cells from 3 experiments. The *x-* axis is discontinued between 0.20 and 0.8 to highlight differences at the cell edges. **k,**, Normalized intensity of TMRM in NHE1-overexpressing cells in 15 kPa PA-based collagen I-coated microchannels in the presence of the mitochondrial calcium uniporter activator kaempferol (Kaemp.) or vehicle control (Ctrl.; DMSO). Data represent mean±S.D. for n=31 cells from 3 experiments. **l,** Arp3 immunofluorescence images of NHE1-overexpressing cells in 15 kPa PA-based collagen I-coated microchannels in the presence of kaempferol or vehicle control (DMSO). Scale bars: 5 µm. **m**, Arp3 intensity normalized to the cell rear and plotted along the normalized cell length in confined NHE1-overexpressing cells in the presence of kaempferol or vehicle control (DMSO). Data are the moving average ± S.D. for n=16 cells from 2 experiments. The *x-* axis is discontinued between 0.1 and 0.8 to highlight differences at the cell edges. **n**, Cell migration velocity of NHE1-overexpressing cells in 15 kPa PA-based microchannels in the presence of kaempferol (Kaemp.) or vehicle control (Ctrl.; DMSO). Data represent mean±S.D. for n=40 cells from 3 experiments. **o**, Number of mCherry wild-type or NHE1-overexpressing cells with or without exogenous mitochondrial addition intravasated to the brain in a zebrafish model 1 day after injection in the perivitelline space (PVS). Data are mean ± S.D. for n≥14 fish from 4 experiments. Statistical significance was assessed by one-way ANOVA followed by Tukey’s (a,c,f) after log transformation (b,o) Mann-Whitney (n), Kruskal-Wallis (i), unpaired t-test (h) after log transformation (k). Cell model: MDA-MB-231.

**Extended Data Figure 8.**
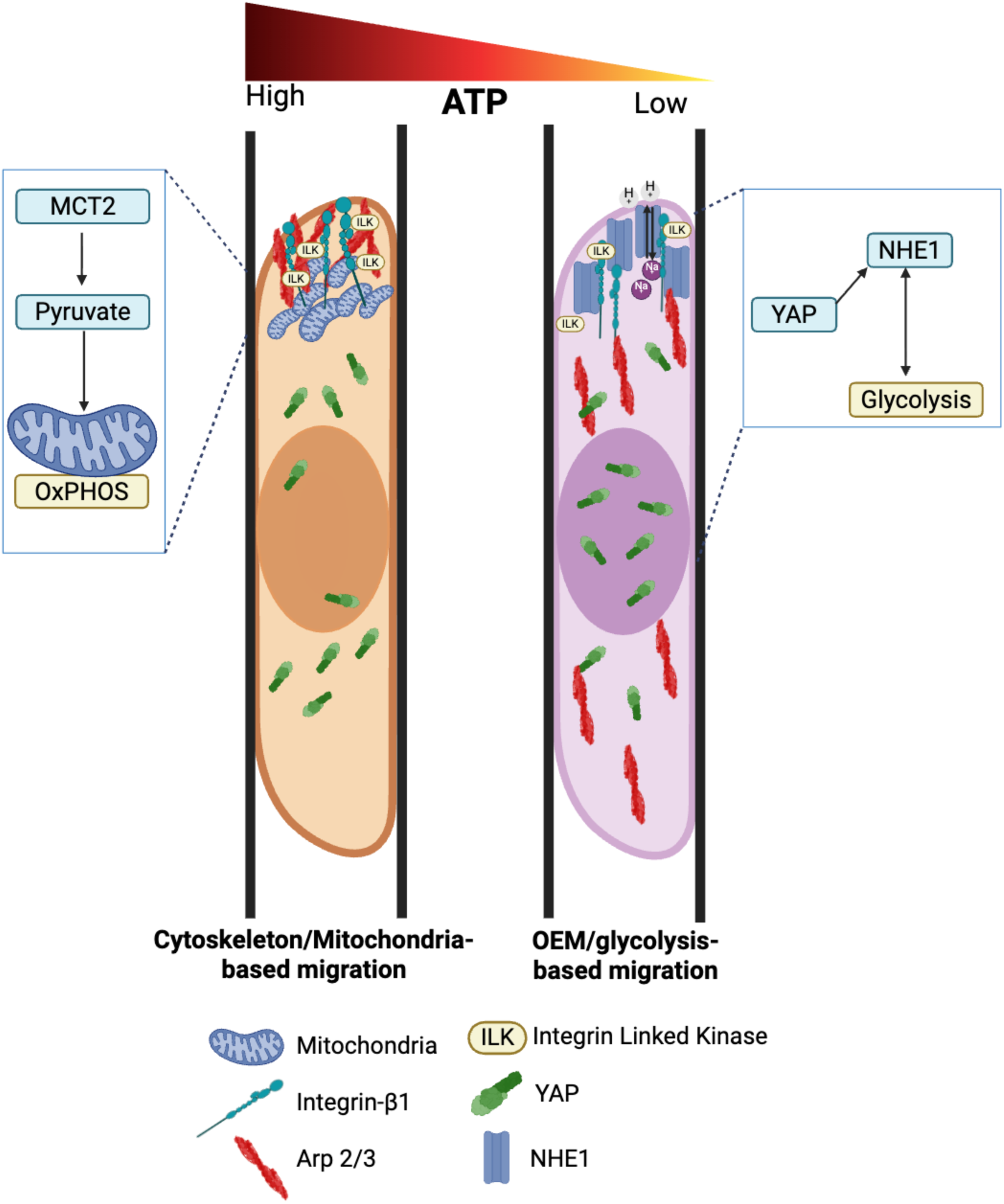
Schematic representing cytoskeleton/mitochondria-dependent versus OEM/glycolysis-dependent confined cell migration.

